# The Siberian wild apple, *Malus baccata* (L.) Borkh., is an additional contributor to the genomes of cultivated European and Chinese apples

**DOI:** 10.1101/2021.09.19.460969

**Authors:** Xilong Chen, Amandine Cornille, Na An, Libo Xing, Juanjuan Ma, Caiping Zhao, Yibin Wang, Mingyu Han, Dong Zhang

## Abstract

It is crucial to understand domestication to unravel the evolutionary processes that shape the divergence of populations. Differences in life-history traits have probably led to marked differences in the mode and speed of evolution between trees and annuals, particularly the extent of crop-wild gene flow during domestication. Apple is an iconic tree and major fruit crop grown worldwide. The contribution of wild apple species to the genetic makeup of the cultivated apple genome remains a topic of intense investigations. We used population genomics in combination with SNPs to investigate the contributions of the two known wild apple relatives, *Malus sylvestris* and *Malus sieversii*, and a supposed contributor, *Malus baccata*, to European and Chinese rootstock and dessert genomes, with a focus on the extent of wild-crop gene flow during apple domestication. We showed that the European dessert and rootstock apples form a specific gene pool, whereas the Chinese dessert and rootstock apples were a mixture of three wild gene pools. Coalescent-based inferences and gene flow estimates indicated that *M. baccata* is an additional contributor to the genome of both European and Chinese cultivated apples through wild-to-crop introgressions. We also confirmed previous results on the contribution of *M. sylvestris* to the cultivated apple genome, and provided insights into the origin of the apple rootstock. This study further demonstrates the role of gene flow during apple domestication, as seen in other woody perennials, and show that domestication of the apple tree involved several wild apple species.

## Introduction

It is crucial to understand domestication to unravel the evolutionary processes that shape the divergence of populations, varieties, and species (Gaut, Díez, & Morrell, 2015; Glémin & Bataillon, 2009; Meyer, DuVal, & Jensen, 2012). Several studies have investigated the evolutionary history of fruit crop perennials by using genome-wide data (Aravanopoulos, Ganopoulos, & Tsaftaris, 2015; Besnard, Terral, & Cornille, 2018; Cao et al., 2016; Chagné et al., 2014; Daccord, Celton, Linsmith, Becker, Choisne, Schijlen, van de Geest, Bianco, Micheletti, Velasco, Di Pierro, Gouzy, Rees, et al., 2017; Harrison & Harrison, 2011; Hazzouri et al., 2015; Huang et al., 2016; Michael & VanBuren, 2015; Teh et al., 2017; Verde et al., 2013). However, documentation of the domestication of fruit tree crops still lags behind that of annual crop plants (Burgarella et al., 2019; Gaut et al., 2015; Meyer et al., 2012; Miller & Gross, 2011; Neale, Martínez-García, De La Torre, Montanari, & Wei, 2017). Differences in life-history traits have probably led to marked differences in the mode and speed of evolution between trees and annuals (Gaut et al., 2015), particularly the extent of gene flow during domestication. Documentation of the domestication of fruit tree crops is important not only academically but also for food production. Because of long-term cultivation, fruit trees are susceptible to ever-changing environments due to climate changes and pathogen outbreaks. Sustainable fruit production is now believed to highly rely on untapped wild genetic diversity (Zhang & Batley, 2020; Zhang, Mittal, Leamy, Barazani, & Song, 2017).

Apple is an iconic tree and major fruit crop grown worldwide. It is also a model species for understanding the role of gene flow during the domestication of clonally propagated perennial crops (Cornille et al., 2019; Cornille, Giraud, Smulders, Roldán-Ruiz, & Gladieux, 2014; Peace et al., 2019; Spengler, 2019). The domestication of apple started in Central Asia, from the local Central crab apple *Malus sieversii* (Ledeb.) M. Roem. Then, apple cultivation spread towards the West, Europe and beyond, and the cultivation was assisted by the possibility of propagating interesting genotypes by grafting (Cornille et al., 2012). Thus, grafting has been an important part of apple evolution and breeding (Wang et al., 2019). With the arrival of the cultivated apple in Europe about 1,500 years ago, the local crab apple *Malus sylvestris* Mill. contributed substantially to the genome of some dessert and cider varieties through wild-to-crop introgressions (Cornille et al., 2012). Population genetic approaches in combination with microsatellite markers have demonstrated that wild-to-crop gene flow has played a major role in the evolutionary history of the cultivated apple in Europe (Cornille et al., 2012). However, while the history of the dessert and cider apples is now well documented in the western part of Eurasia (Bina et al., 2021; Cornille et al., 2014; Migicovsky et al., 2021; Peace et al., 2019; Spengler, 2019), the domestication history of apples in the East remains unclear. In addition, the history of the apple rootstock that allowed the spread of apple across the world needs to be elucidated. Understanding the origin of the apple rootstock can also provide insights into the role of crop-wild gene flow, as several rootstocks are supposed to be the result of wild-crop hybridization and are often dwarf forms of local wild species (Volk, Cornille, Durel, & Gutierrez, 2021; Wang et al., 2019).

In China, apple cultivation has a tangled and complex history that spans a millennium and involves several wild and cultivated apple species, used for dessert or as rootstocks, that were crossed by ancient communities or bred recently. For more than 2,000 years, ancient Asian communities cultivated *Malus × domestica* ssp. *chinensis* Li Y.N. (called Mianpingguo, which means soft apple) (Gao et al., 2015; Yunong, 1999). *Malus × domestica* ssp. *chinensis* is supposed to originate from *M. sieversii* located in the northeastern part of China (Xinjiang Province), close to the borders of Kazakhstan and Kyrgyzstan. Then, the cultivated species was propagated along the Silk Routes across China where several local cultivars once existed (Gao et al., 2015). However, from the 18th century, most of the local cultivars disappeared, and, nowadays, only a few Chinese landraces remain, namely, Mianpingguo, Shaguo, Huahong, and Caiping. These four Chinese cultivars are supposed to belong to two species: *M. domestica* subsp. *chinensis* and *Malus asiatica*. Some hybrids of *M. domestica* ssp. *chinensis* and *M. asiatica* are cultivated in China. In addition, some modern apple varieties from Western countries, such as Golden Delicious, Gala, and Fuji, are also cultivated in China, and they were introduced from Europe and the United States less than a decade ago. A few other cultivated Chinese apple species are traditionally used as rootstocks and for ornamental purposes, and some of them originated from several hybridization events (Gao et al., 2015). *Malus prunifolia* (Chinese crabapple called Qiuzi) and *Malus robusta* (*i.e.*, *Malus baccata* × *M. prunifolia* cross called Balenghaitang) are used as rootstocks. *Malus micromalus*, *Malus halliana*, and *Malus spectabilis* are used as ornamental species. These rootstocks and ornamental species cannot be found in natural habitats in China, so they are classified as cultivated species (Yunong, 1999). The Chinese people have also used local wild apple species for millennia. Because of their ease of propagation through apomixis and their cold resistance, *M. baccata* and *Malus hupehensis* are supposed to be the progenitors of the Chinese rootstocks. Indeed, *Malus baccata*, the Siberian wild apple, is the main genetic resource (especially as a rootstock) for apple breeding programs in China, because of its excellent resistance to cold stress and apple scab (Chen et al., 2019; Gygax et al., 2004; Volk et al., 2015). *Malus hupehensis*, the tea crabapple or Hupeh crab, is an apomictic species. Its seedling rootstocks can be propagated directly through seedlings, instead of using cuttings or layering propagation (Wang et al., 2019; Yang, Duan, & Zhang, 2008). China also introduced European-bred dwarf rootstocks in the 19th century, but most apple production areas in China still use local cultivated (*M. prunifolia* and *M. robusta)* and wild (*M. hupehensis* and *M. baccata*) apples as rootstocks because of their good adaptability (Wang et al., 2019).

Apple domestication and breeding in China have therefore probably involved crop-crop and wild-crop gene flow and contributions of several wild species. However, all available information on apple domestication in China is derived from historical records and traditional knowledge. Only Duan et al. (2017) showed that *M. asiatica* and *M. prunifolia* originated from crosses between *M. sieversii* and *M. baccata*, but this result remains anecdotic. It is therefore still unclear whether cultivated Chinese dessert and rootstock apples originated from *M. sieversii* and/or *M. baccata* or/and *M. hupehensis,* or both originated from *M. sieversii* followed by recurrent wild-to-crop gene flow from *M. baccata* or/and *M. hupehensis*. Besides, the genetic relationships among wild and cultivated European and Chinese apples have not yet been elucidated, and it is unclear whether substantial crop-to-wild gene flow occurred during apple domestication in China, as observed in Europe and the Caucasus (Cornille et al., 2012; Bina et al., 2021).

Here, we investigated the history of domestication of European and Chinese dessert apples and rootstocks, with a focus on the extent of crop-wild gene flow during domestication. We combined previously published sequenced genomes (Duan *et al*., 2017) with a new dataset that included wild and cultivated Chinese apples sequenced for their RNA. Using single-nucleotide polymorphisms (SNPs) called from several wild apple species, including the wild apple relatives of *M. domestica* (*i.e.*, *M. sieversii* and *M. sylvestris*), *M. baccata*, *M. hupehensis*, and other ancestral species from the genus *Malus*, as well as European and Chinese apple cultivars, we aimed to answer the following questions: (1) What is the population structure and differentiation of wild and cultivated apples from Eurasia? (2) Did gene flow occur between wild and cultivated apples during domestication? (3) What is the domestication history of Chinese and European dessert apple and rootstock cultivars and were they independently domesticated from different wild species?

## Materials and Methods

### Plant samples

For this study, fruit flesh samples of 71 *Malus* accessions, representing 48 wild and 23 cultivated apple samples, were collected from repositories in China (Table S1). Additionally, short-read DNA sequences of 97 *Malus* accessions (Duan *et al*., 2017), namely, 53 wild apple samples and 44 apple cultivars, were downloaded (SRA code accession: SRP075497, Table S2). The two datasets, which included 168 accessions with 66 apple cultivars (7 historical Chinese cultivars, 30 cultivars from Europe or Western countries, 25 rootstocks, and 4 ornamental cultivars) and 102 wild apples (43 *M. sieversii*, 18 *M. baccata*, 11 *M. sylvestris*, and 30 other *Malus* accessions), were merged for subsequent analyses (Table S1).

### RNA extraction and library construction and sequencing

The fruit flesh of the 71 collected samples was immediately transferred to liquid nitrogen and stored at -80°C until RNA extraction. The RNA was extracted using the SDS–phenol method (Hu et al., 2002). The RNA quality was checked using an agarose gel, and RNA concentrations were estimated with a NanoDrop 1000 spectrophotometer (NanoDrop Technologies, Wilmington, DE, USA). After RNA extraction and DNase I treatment, mRNA was isolated from the total RNA by using magnetic oligo (dT) beads. The mRNA was mixed with fragmentation buffer and cleaved into short fragments and used as templates for cDNA synthesis. The short fragments were purified, resolved with EB buffer for end reparation and single adenine nucleotide addition and connected with adapters. After agarose gel electrophoresis, suitable fragments were selected as templates for polymerase chain reaction (PCR) amplification. During the quality control steps, an Agilent 2100 Bioanalyzer and ABI StepOnePlus Real-Time PCR system (Agilent, US) were used for evaluating the quantity and quality of the sample libraries. Finally, the constructed libraries were sequenced on an Illumina HiSeq 2000 system (BGI, Shenzhen, China). Quality control of the raw data was performed with FastQC (http://www.bioinformatics.babraham.ac.uk/projects/fastqc), and 420.73 GB of clean data was obtained (Table S3).

### Read mapping and SNP calling and filtering

Two different pipelines were used for the sequenced DNA and RNA reads (Figure S1). For the sequenced RNA accessions, reads were processed as follows: SNP calling was performed on the basis of the transcript sequence data with GATK v3.5 calling variants pipeline for RNAseq (McKenna et al., 2010) (https://gatk.broadinstitute.org/hc/en-us/articles/360035531192?id=3891). Filtered reads were mapped to the new high-quality *M. domestica* reference genome GDDH13 v1.1 (Daccord *et al*., 2017) with Hisat2 software (Kim, Langmead, & Salzberg, 2015). The SAM file was processed with Picard (*Picard toolkit*, 2019) in several steps, as described by ForgeMIA. The GATK module HaplotypeCaller was used for SNP calling, and low-quality SNPs were filtered out with the GATK module VariantFiltration. For short-read DNA sequences of the 97 accessions (Duan *et al*., 2017), SNPs were called using the GATK v3.5 pipeline (McKenna et al., 2010). The reads were aligned to the GDHH13 v1.1 *M*. *domestica* reference genome (Daccord *et al*., 2017) with BWA by using the “mem” algorithm (Li, 2013). The redundant reads were removed using the MarkDuplicate module from Picard (“Picard Toolkit,” 2019). The GATK module HaplotypeCaller was used for SNP calling, and GenotypeGVCFs module was used to produce a raw set of joint SNPs. The raw joint SNPs were finally filtered with the GATK VariantFiltration module and default quality thresholds.

### Suitability of the markers used for phylogenetic, population structure, and demographic inferences

The filtering steps used for the analyses described below are provided in Figure S1. The SNP files obtained from the RNA-seq and DNA-seq datasets were combined with the VCFtools vcf-merge function (Danecek et al., 2011). The clone and duplicate samples were removed (*i.e.*, individuals with pairwise KING-robust kinship estimates among individuals > 0.354) (Manichaikul et al., 2010)) with Plink 2.0 (Chang et al., 2015). The SNPs with minimum minor allele frequency <□0.01 and minimum site coverage = □0.9 were filtered out with Plink (Purcell et al., 2007). A total of 76,239 SNPs were obtained. For population structure and demographic inferences, linked and non-synonymous SNPs were filtered out to avoid any bias associated with potential SNPs linked to the selected genomic region. First, SNPs in linkage disequilibrium (LD) were removed; the LD for apples was <1,000 bp (Duan *et al*., 2017). The linked SNPs were filtered out with Plink (Purcell *et al*., 2007) by using the following parameters: window = 1 kb, step = 1 SNP, and *r^2^* = 0.2. Second, the remaining SNPs were annotated with SnpEff 4.0 (https://pcingola.github.io/SnpEff/) (Cingolani et al., 2012), and only synonymous SNPs were retained; therefore, 36,200 unlinked and synonymous SNPs were used (Figure S1).

### Population structure, genetic diversity, and differentiation among wild and cultivated apples

The population structure and admixture among *Malus* accessions were inferred using a model-based clustering method implemented in ADMIXTURE v.1.23 (Alexander, Novembre, & Lange, 2009). ADMIXTURE is based on the same statistical model as the one implemented in STRUCTURE (Pritchard, Stephens, & Donnelly, 2000), but it uses a fast numerical optimization algorithm for large SNP datasets to infer the proportion of ancestry of genotypes in *K* distinct predefined clusters. The number of ancestral populations, *K*, was varied from 1 to 10, with 20 repetitions per *K* and default settings. The consensus solution for each *K* value was generated with CLUMPAK (Kopelman, Mayzel, Jakobsson, Rosenberg, & Mayrose, 2015). The amount of additional information explained by increasing *K* was determined using the cross-validation procedure of ADMIXTURE (Alexander et al., 2009). However, the *K* identified with the cross-validation procedure often does not correspond to the finest biologically relevant population structure (Cornille et al., 2015; Kalinowski, 2011; Puechmaille, 2016). Therefore, the bar plots were visually checked, and the *K* value for which all clusters had well-assigned individuals was selected (*i.e.*, that no further well-delimited and relevant clusters could be identified for higher *K* values than the chosen one). All the bar plots were generated with the R package “Pophelper” (Francis, 2017)

The genetic variation and differentiation among the groups detected with ADMIXTURE were further investigated using three different methods. First, a principal component analysis (PCA) was performed with the dudi.pca function of the ade4 package in R (Dray & Dufour, 2007). The first two principal components were plotted with the R package “ggplot2” (Wickham, 2009). Second, a neighbor-net tree based on Nei’s distance (Masatoshi Nei, 1978) among individuals was created. The neighbor net was visualized with Splitstree v4 (Huson & Bryant, 2006). For the PCA and neighbor-net tree, individuals assigned to a given cluster with a membership coefficient ≥ 0.55 were indicated using the color of each cluster, and admixed individuals (*i.e.*, individuals with a membership coefficient to any given cluster < 0.55) were colored gray. We chose this threshold because both cultivated and wild apples were highly admixed, and the use of a higher cut-off would have removed too many individuals (see Results). For each wild or cultivated population (*i.e.*, group of individuals with membership coefficient > 0.55 to a given cluster detected with ADMIXTURE), Nei’s diversity (π) (M Nei & Li, 1979), observed (*H_O_*) and expected heterozygosity (*H_E_*) (Masatoshi Nei, 1973), inbreeding coefficient *F_IS_*, and genetic differentiation (*F_ST_*) between populations were estimated with Stacks population program v2.52 (Rochette, Rivera-Colón, & Catchen, 2019).

### Detection of gene flow

Two approaches were used to investigate the occurrence of gene flow among the populations (*i.e.*, group of individuals with a membership coefficient > 0.55 to a given genetic cluster detected with ADMIXTURE).

First, TreeMix (Pickrell & Pritchard, 2012) with the ipyrad tool (Eaton & Overcast, 2020) was used to analyze the relationships among the populations and the potential influence of gene flow. TreeMix uses genome-wide allele frequency data to estimate the maximum likelihood tree of populations. Then, on the basis of populations with the poorest fit to the tree model, the program infers the presence and magnitude of migration events between them (Pickrell & Pritchard, 2012). Trees were rooted with the *Malus* species from North America, which are known to be the most divergent and ancestral *Malus* group (Harris *et al*., 2002 and see Results). Zero to 9 migration numbers were tested, and the run for which the likelihood of trees reached an initial plateau (Aguirre-Liguori et al., 2019; Brandenburg et al., 2017) and the magnitude of migration rates decreased was retained.

Second, Dtrios of Dsuite (Malinsky, Matschiner, & Svardal, 2021) was used to perform four-taxon *D-statistic* tests on the final unlinked synonymous SNP dataset. *D-statistics* is also called Patterson’s *D* (ABBA-BABA) statistic (Durand, Patterson, Reich, & Slatkin, 2011; Green et al., 2010), and it checks whether gene flow occurred among populations (i.e., group of individuals assigned with a membership coefficient > 0.55 to a given genetic group) with ADMIXTURE and 36,200 synonymous unlinked SNPs. Dsuite calculates *D-statistics* not only for one specific quartet but also, comprehensively, for sets of populations in a VCF file and keeps the outgroup fixed (here the North American wild apples, see Results). The significance of *D-statistics* was assessed using Jackknife (Green et al., 2010) on 20 blocks. Results for which the Z-score was higher than 3 and *P-value* < 0.01 were retained.

### Approximate Bayesian computation

The history of apple domestication was reconstructed using the approximate Bayesian computation (ABC) framework in combination with a coalescent-based inference simulator, fastsimcoal2 (Excoffier, Dupanloup, Huerta-Sánchez, Sousa, & Foll, 2013). Our aims were to infer (1) whether gene flow occurred among certain crop and wild populations during apple domestication and (2) the European and Chinese cultivated apples originated from which wild apple population(s). Previous studies ( Cornille, Giraud, Smulders, Roldán-Ruiz, & Gladieux, 2014; Cornille et al., 2012; Duan et al., 2017; Migicovsky, Gardner, Richards, Thomas Chao, et al., 2021) have reported that the European dessert apple (*M. domestica*) diverged from *M. sieversii*. Therefore, the origin of the European dessert apple was not tested, and it was assumed that it diverged from *M. sieversii* in our models. The scenarios were established according to the results obtained from ADMIXTURE, PCA, and neighbor-net tree as well as the results of *D-statistics* and TreeMix. The populations were defined as those detected with ADMIXTURE analyses (*i.e.*, group of individuals assigned with a membership coefficient > 0.55 to a given genetic group).

A newly developed ABC method was used for model selection and parameter estimation according to a machine learning tool called “Random Forest” (ABC-RF). This approach allowed us to disentangle complex demographic models (Pudlo et al., 2016) by comparing groups of scenarios with a specific type of evolutionary event to other groups with different types of evolutionary events (instead of considering all scenarios separately) (Estoup, Raynal, Verdu, & Marin, 2018) in what we will hereafter call “ABC rounds.” Such a grouping approach in the scenario choice is more powerful than testing all scenarios individually to disentangle the main evolutionary events that characterize speciation (Estoup et al., 2018).

Two nested sets of ABC analyses were run, and, within each set, several rounds were run (Figure S2). Such a nested ABC approach avoids the comparison of too complex models with numerous populations and parameters (Estoup et al., 2018). The first ABC set tested the history of domestication of the European apple rootstock (set 1, Figure S2). Once the history of the cultivated European apples was inferred, a second set of ABC analyses was run to infer the domestication history of the cultivated Chinese dessert and rootstock apples (set 2, Figure S3). For the two ABC sets, the modalities of gene flow among crop and wild populations were defined based on the TreeMix analyses. Prior distributions for each parameter are provided in Table S4.

For all models, 36,200 unlinked SNPs were simulated, with 8,000 simulations per scenario. For each simulation, the following summary statistics were computed with arlsumstats v 3.5 (Excoffier & Lischer, 2010): the number of sites with segregating substitutions of population i (*S_i*, i = {1, 2, 3, 4,5}), mean number of pairwise differences of population i (π*_i*, i = {1, 2, 3, 4,5}), mean number of differences between pairs of populations (π*_i_j*, i, j = {1, 2, 3, 4,5}, i ≠ j), and pairwise *F_ST_* (*F_ST__i_j*, i, j = {1, 2, 3, 4,5}, i ≠ j). Summary statistics were also added on the basis of the joint frequency spectrum between all pairs of populations (Tellier et al., 2011; Wakeley & Hey, 1997) computed with a home-made script: sites polymorphic in population i, but monomorphic in population j, and vice-versa (*S_x1_i_j_* and *S_x2_i_j_*, respectively); number of shared polymorphic sites between population i and population j (*S_si_*__j_); and number of sites showing fixed differences between populations i and j (*S_f_i_j_*).

The *abcrf* v.1.7.0 R statistical package (Pudlo et al., 2016) was used to perform the ABC-RF analysis, which provides a classification vote that represents the number of times a scenario is selected as the best one among *n* trees in the constructed random forest. For each ABC set, the scenario, or a group of scenarios, with the highest number of classification votes was selected as the best scenario, or best group of scenarios, among a total of 500 classification trees (Breiman, 2001). The posterior probabilities and prior error rates (*i.e.,* the probability of choosing a wrong group of scenarios when drawing model index and parameter values from the priors of the best scenario) over 10 replicate analyses (Estoup et al., 2018) were computed for each ABC step. The simulated models were also visually checked to be compatible with the observed dataset by projecting the simulated and observed datasets onto the first two linear discriminant analysis (LDA) axes (Pudlo et al., 2016) and whether the observed dataset was within the clouds of simulated datasets. The parameter inferences were then calculated using the final selected model, according to the three-set ABC procedure. The ABC-RF approach includes the model checking step performed *a posteriori* in previous ABC methods.

Once the most likely apple domestication history was obtained, posterior distributions were estimated for each parameter and 1,000 pseudo-observed datasets were re-simulated using the 95% confidence interval of the posterior estimates to evaluate the goodness of fit of the final chosen demographic model with *abc* R package (Csilléry, François, & Blum, 2012).

## Results

### The European and Chinese cultivars have different ancestries

We removed 20 individuals of the initial dataset (*N* = 168) because they had unknown passport information (*N* = 1) or were detected as a clone or duplicate (*N* = 19, Table S1). The first run of ADMIXTURE, in which the 20 individuals were removed (*N* = 148), revealed 20 wild apple trees described as *M. sieversii* or *M. sylvestris* or *M. baccata* admixed with the cultivated gene pool (Figures S4 and S5) or misidentified on the field (*i.e.*, fully assigned to *M. domestica* gene pool), as previously observed (Cornille, Gladieux, & Giraud, 2013; Cornille et al., 2015, 2014; Feurtey, Cornille, Shykoff, Snirc, & Giraud, 2017). We also detected 13 wild individuals, described as not one of the three wild apple species studied here, admixed with multiple gene pools (Table S1, Figures S4 and S5). In this study, we focused on the history of the cultivated apple, so we wanted “pure” wild apple reference samples of the three wild apple relatives, *i.e.*, *M. baccata*, *M. sieversii*, and *M. sylvestris*. Therefore, we filtered out the 33 wild admixed or misidentified individuals detected with the first ADMIXTURE analysis (Figure S6, Table 1). We retained *M. hupehensis*, as the samples belonging to this species grouped with *M. baccata* at a high membership (Figure S6). Population genetic structure inferences for the whole dataset (*N* = 148) vs. the pruned dataset for those admixed individuals (*N* = 115) were consistent (Figures S4 and 1a). However, ADMIXTURE revealed a clearer population genetic structure for the dataset pruned for wild admixed individuals. Therefore, we retained the dataset pruned for wild admixed or misidentified individuals for further analyses.

**Table 1.**
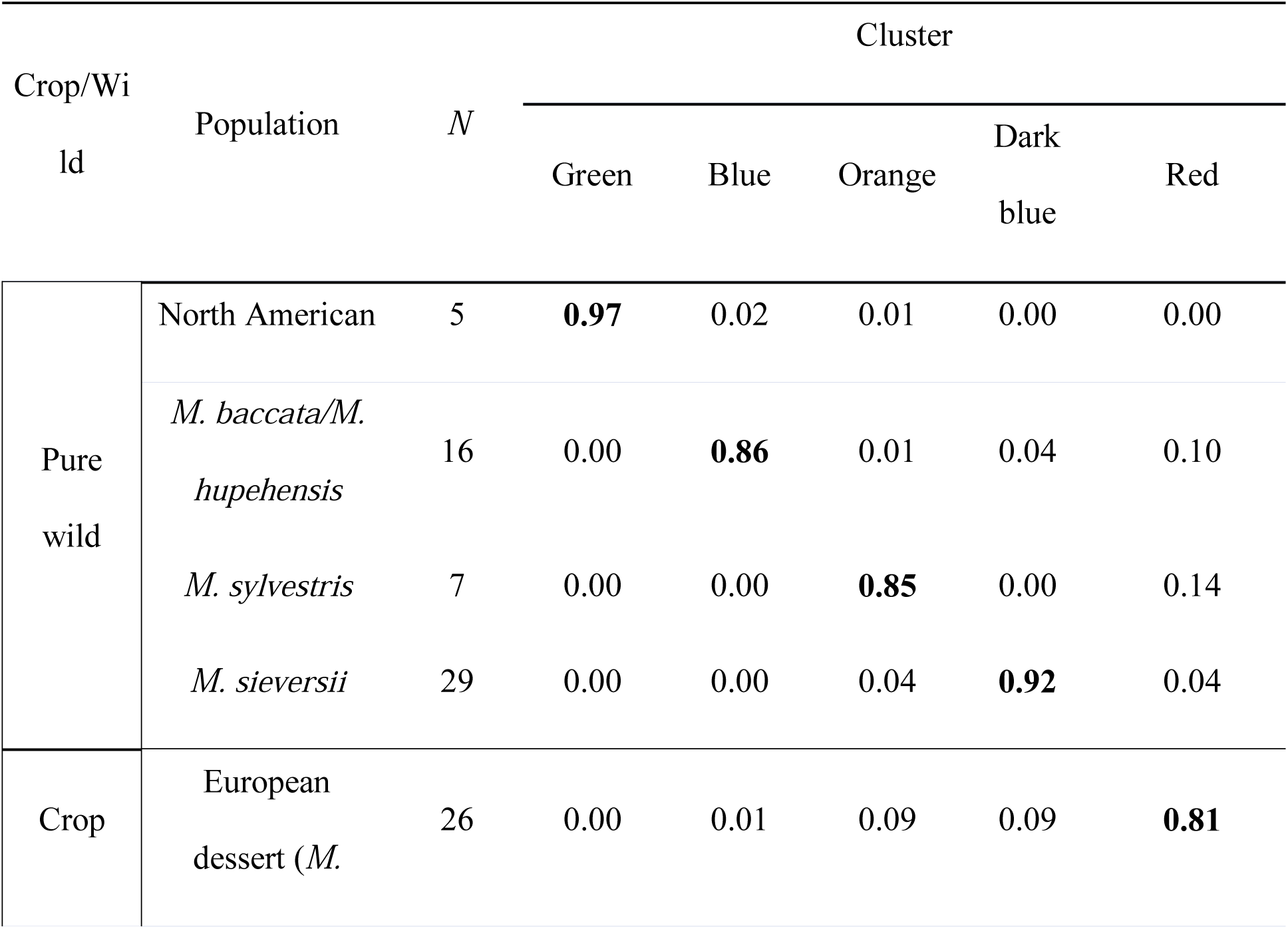
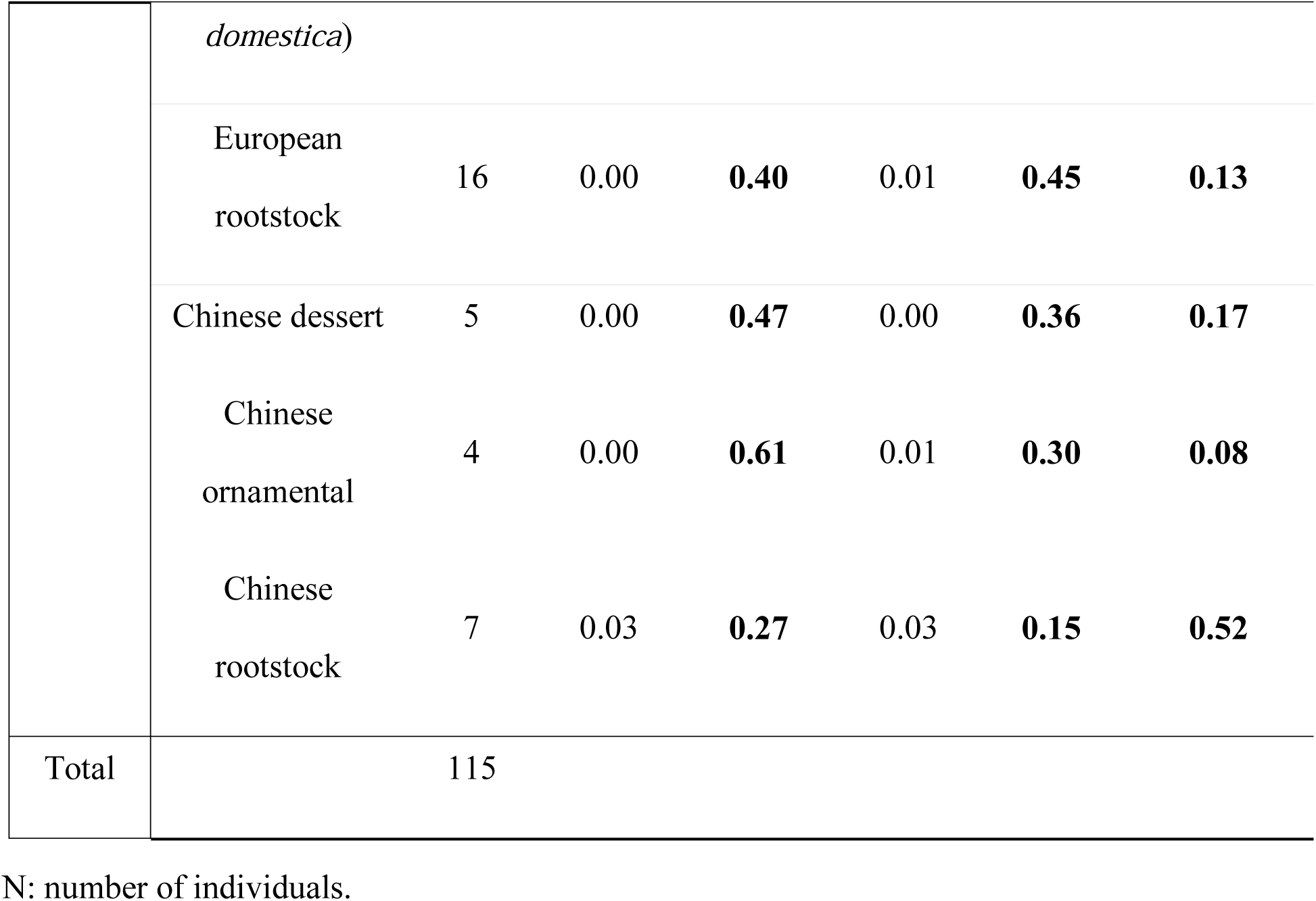
Mean proportions of nine cultivated and wild apple populations assigned to five genetic clusters inferred with ADMIXTURE at *K* = 5 (*N* = 115, 36,200 SNPs). We divided the cultivated apple samples into five populations according to ADMIXTURE inferences and their origin (Europe or China) and use (dessert or rootstock). We assigned only wild populations with membership coefficient > 0.55 to a given genetic group. Thus, samples were partitioned into nine populations, namely, four wild and five cultivated apple populations.

ADMIXTURE revealed five main genetic groups (Figure 1a). Cross-validation error monotonically decreased up to *K* = 8 (Figures S7 and S8). At *K* > 5, further substructure was observed, but new clusters were represented by only admixed individuals (Figure S7). Besides, at *K* = 5, the population structure was consistent with the morphological classification of the wild species ( Cornille et al., 2019, 2014, 2012; Harris et al., 2002; Robinson, Harris, & Juniper, 2001), with *M. sylvestris* (orange), *M. sieversii* (purple), *M. baccata/M. hupehensis* (blue), and North American wild apples (green) each forming a distinct group. At *K* = 5, the European apple cultivars (*i.e.*, *M. domestica*, including European dessert and rootstock cultivars) formed a distinct genetic group highly admixed with *M. sylvestris* (Table 1), as shown by Cornille et al. (2012), but also with the *M. baccata/M. hupehensis* (blue) genetic group (Table 1). For *K* = 5, the Chinese dessert and rootstocks as well as ornamental Chinese apple species did not form a specific genetic group, but a genetic mixture of the blue (*M. baccata/M. hupehensis*) and purple (*M. sieversii*) gene pools (Table 1) as well as, to a lesser extent, with the orange genetic group (*M. sylvestris*).

**Figure 1.**
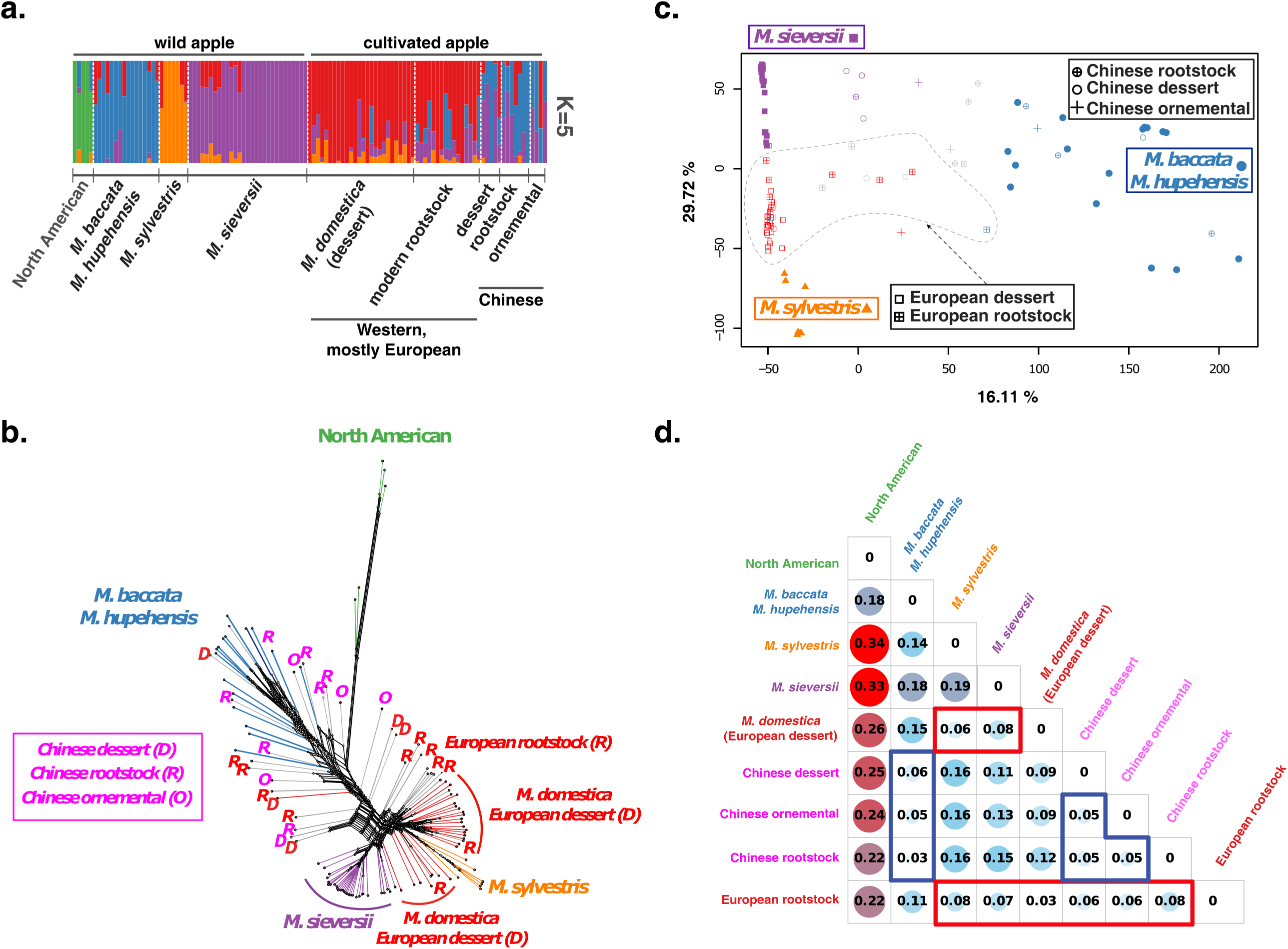
Genetic structure, variation, differentiation, and relationships of wild and cultivated European and Chinese apples. a. Population structure inferred with ADMIXTURE at *K* = 5 (*N* = 115, 36,200 unlinked synonymous SNPs); each individual is represented by a vertical bar partitioned into *K* segments that represent the proportions of ancestry of its genome in *K* clusters. Colors represent the inferred ancestry from *K* ancestral genetic clusters. b. Neighbor net tree depicting the relationships among the wild and cultivated apple populations identified with ADMIXTURE for *K* = 5. Colors correspond to the genetic groups inferred at *K* = 5, and admixed samples (*i.e.*, individuals with membership coefficient < 0.55 to any given gene pool) are in grey. c. Principal component analysis (PCA) representing the genetic variation among wild and cultivated apples, with the total variance explained by the first two components (29.72% and 16.11%, respectively). d. Pairwise genetic differentiation (*F_ST_*) among populations identified with ADMIXTURE at *K* = 5 (excluding wild individuals with membership coefficient < 0.55 to a given gene pool).

Then, we assessed the genetic variations among wild and cultivated apple populations detected with ADMIXTURE. We assigned wild individuals with a membership coefficient > 0.55 to a given cluster in the corresponding population (Table 1). Because of the high level of admixture in cultivated apples, we retained all cultivars and defined five cultivated apple groups based on the ADMIXTURE results as well as their historical uses and geographic origins (Tables 1 and S1). The use and geographic origins of the cultivars were documented on the basis of the studies performed by Morgan & Richards (1993, 2003) and Urrestarazu *et al*. (2016) for the European-Western cultivars and Zhi-Qin (1999) for the Chinese cultivars. The North American and Siberian wild apples were the most genetically differentiated population (Figure 1b, c, d). *Malus sieversii* individuals formed a distinct group (Figure 1b), close to that of the European dessert apple. The European cultivars (*M. domestica*) were closer to *M. sylvestris* than to the known progenitor *M. sieversii* (Figure 1b, c, d), as detected previously ( Cornille et al., 2012; Duan et al., 2017; Migicovsky, Gardner, Richards, Thomas Chao, et al., 2021). *Malus sylvestris* was nested with *M. domestica* in the neighbor-net tree (Figure 1b), but it formed a distinct, but close, gene pool to that of *M. domestica* in the PCA (Figure 1c). Most of the European rootstocks were admixed with the three wild apple species (Figure 1b, c). The Chinese dessert and rootstock formed a bushy structure intermingled with *M. baccata* and *M. hupehensis*. *F_ST_* estimates further confirmed that the European cultivars were genetically close to *M. sylvestris* and *M. sieversii*, and the Chinese cultivars were genetically close to *M. baccata-M. hupehensis* (Figure 1d). Genetic diversity estimates for the nine populations are provided in Figure S9 and Table S5. Genetic diversity was significantly lower in the wild apple than in the cultivated apple gene pools, except for the *M. baccata/M. hupehensis* group; this can be explained by the high level of admixed individuals in the cultivated populations (Table 1, Figure 1).

Therefore, ADMIXTURE, neighbor-net tree, PCA, and *F_ST_* suggest that the cultivated European and Chinese apples have different ancestries, which raise questions about their origin and whether crop-wild gene flow occurred. The European dessert and rootstock cultivars grouped together (*M. domestica*) and formed a specific gene pool; this has been previously reported for the dessert apples ( Cornille et al., 2012; Migicovsky, Gardner, Richards, Thomas Chao, et al., 2021). The European dessert and rootstock cultivars showed different levels of admixture with the wild species: the European dessert cultivars were mostly admixed with *M. sylvestris* (and *M. sylvestris* was nested in *M. domestica*), whereas the European rootstock cultivars were mainly admixed with *M. baccata* and *M. sieversii*. The high level of admixture of the European rootstock with *M. baccata* and *M. sieversii* raises questions about the origin of the European rootstock cultivars: their distinct genetic differentiation from *M. baccata* and clustering with *M. domestica* (which we know originated from *M. sieversii*) may suggest that *M. baccata* contributed to the European rootstock through recent crop-to-wild introgressions and the European rootstocks originated from *M. sieversii*. In contrast to the European cultivars (*M*. *domestica*), the Chinese cultivars, both dessert and rootstock, did not form a specific gene pool, but they were a mixture of *M. sieversii* or *M. baccata* gene pools. The high level of admixture of the Chinese cultivars (rootstock or dessert) with *M. baccata* and *M. sieversii* and even, sometimes, their full membership to genetic clusters of two wild apple gene pools, suggests that wild apple trees are grown in orchards for consumption without any strong domestication process in China and/or substantial wild-crop gene flow occurred. In addition, the observation that the Chinese cultivars are nested and genetically close to *M. baccata* (in contrast, *M. domestica* was grouped with *M. sieversii* and *M. sylvestris*) suggests that *M. baccata* may be the wild progenitor of the Chinese cultivars; domesticated populations are expected to be nested within their source population because they recently diverged from a subset of individuals within the source population (Matsuoka et al., 2002). Although our analyses showed that *M. sylvestris* is nested within *M. domestica*, *M. sylvestris* is known to not be a progenitor of the cultivated apple; it is a secondary contributor through crop-to-wild introgressions. Therefore, we investigated the contributions of each wild species by crop-to-wild gene flow or/and an initial domestication event.

### Substantial wild-to-crop gene flow from the three wild apple relatives

TreeMix (Figures 2 and S10) indicated the occurrence of gene flow from the European wild apple to the European dessert and rootstock, but not from *M. sieversii*. TreeMix also indicated gene flow from *M. sieversii* to the Chinese apple rootstock and from *M. baccata* to the Chinese dessert. The D-suite inferences were congruent with TreeMix results, with significant excess of sharing of derived alleles among almost all crop and wild population pairs (Table S6). In addition, TreeMix analyses suggested that the cultivated European apples formed a monophyletic group close to *M. sieversii*; in contrast, the Chinese cultivars formed a polyphyletic group, with the Chinese dessert apple being a sister group with *M. sieversii* and Chinese rootstock, a sister group with *M. baccata*. However, TreeMix can be biased by its starting tree, especially when many populations are admixed (Lipson, 2020; Lipson et al., 2013). Therefore, we investigated the domestication history of the Chinese and European cultivars by using coalescent-based approaches, which allow us to disentangle between crop-wild gene flow and ancestral polymorphisms.

**Figure 2.**
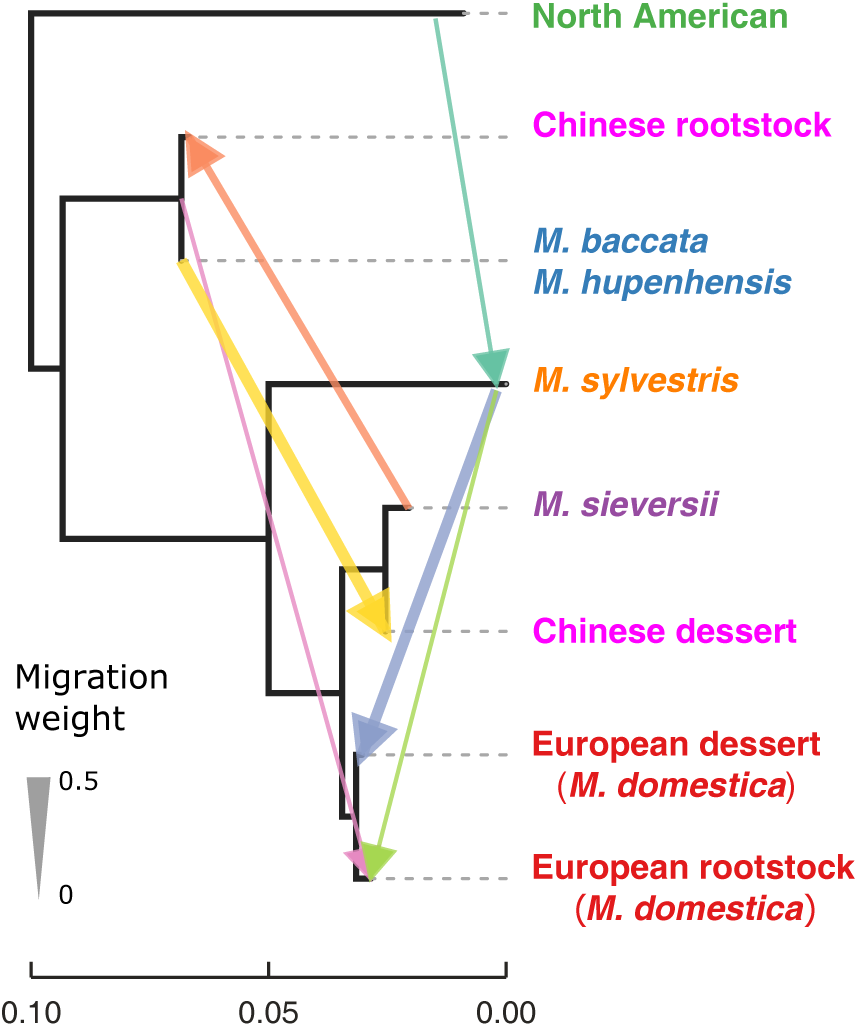
Gene flow inferred with TreeMix (*N* = 114, 36,200 SNPs; Chinese ornamentals were removed because of low number of individuals, *N* = 4) among the wild and cultivated apple populations detected with ADMIXTURE at *K* = 5. The North American wild apples were used as the outgroup. The best number of migration events was six (Figure S10). Arrows of different colors represent migration events.

### Domestication of the European and Chinese apples from *M. sieversii-M. domestica* followed by substantial gene flow from the Siberian and European wild apples

For ABC inferences, we excluded the North American wild apple population, as it was the most genetically distant and did not contribute to the cultivated apple genome in the analyses (Figure 1). We also excluded the Chinese ornamental apple because it included only four individuals. Our aim was to analyze (1) the extent of crop-wild gene flow and (2) infer the divergence history of the cultivated apple populations (excluding the European dessert, as its history is already known) (Cornille et al., 2012; Duan et al., 2017; Harris et al., 2002; Migicovsky, Gardner, Richards, Thomas Chao, et al., 2021; Sun et al., 2020; Velasco et al., 2010).

First, we inferred the origin of the cultivated European rootstock apples (Figures S2 and 3) with respect to the three wild apple populations and European dessert. We assumed that three wild apples diverged from an unknown ancestral population and the European dessert apple *M. domestica* diverged from *M. sieversii*. Our results above (population structure and gene flow estimates) suggest that *M. baccata-M. hupehensis* did contribute to the European rootstock through recent crop-to-wild introgressions and the European rootstock cultivars originated from *M. sieversii*. For the first set of ABC (Figure S2), we tested whether the European rootstock diverged from *M. domestica* or *M. sieversii* (Figure S2) and assumed four different modalities of gene flow: no gene flow, gene flow between European dessert and *M. sylvestris* only (Cornille et al., 2012), gene flow between the European rootstock and *M. sylvestris* and *M. baccata/M. hupehensis* only, and a combination of the first two modalities of gene flow (Figure S2). The last modality was defined on the basis of an additional TreeMix analysis that focused on only the populations used for ABC set 1 (Figure S11). Therefore, we simulated eight scenarios for the first ABC set (Figure S2). For the second set of ABC analysis (set 2, Figure S3), we inferred the domestication history of the Chinese dessert and rootstock apple on the basis of the most likely group of scenarios selected from the first ABC set. We tested whether the Chinese desert and rootstock cultivated apples diverged from (i) *M. domestica*, (ii) *M. baccata-M. hupehensis*, or (iii) *M. sieversii*. We defined 12 different scenarios of divergence of the Chinese cultivars and assumed two modalities of gene flow: gene flow as inferred from TreeMix (Figure 2) and no gene flow between the Chinese rootstock and *M. baccata-M. hupehensis* and *M. sieversii +* no gene flow between the Chinese dessert with *M. baccata-M. hupehensis* and *M. sieversii* (Figure S3). In total, we simulated 24 scenarios. For each ABC round, the observed summary statistics were within the cloud of simulated summary statistics, which did not overlap across the model groups (Figures S9, S10) and indicated that these tests could discriminate among various competing scenarios (except for the fourth round of the second ABC set, Figure 3).

**Figure 3.**
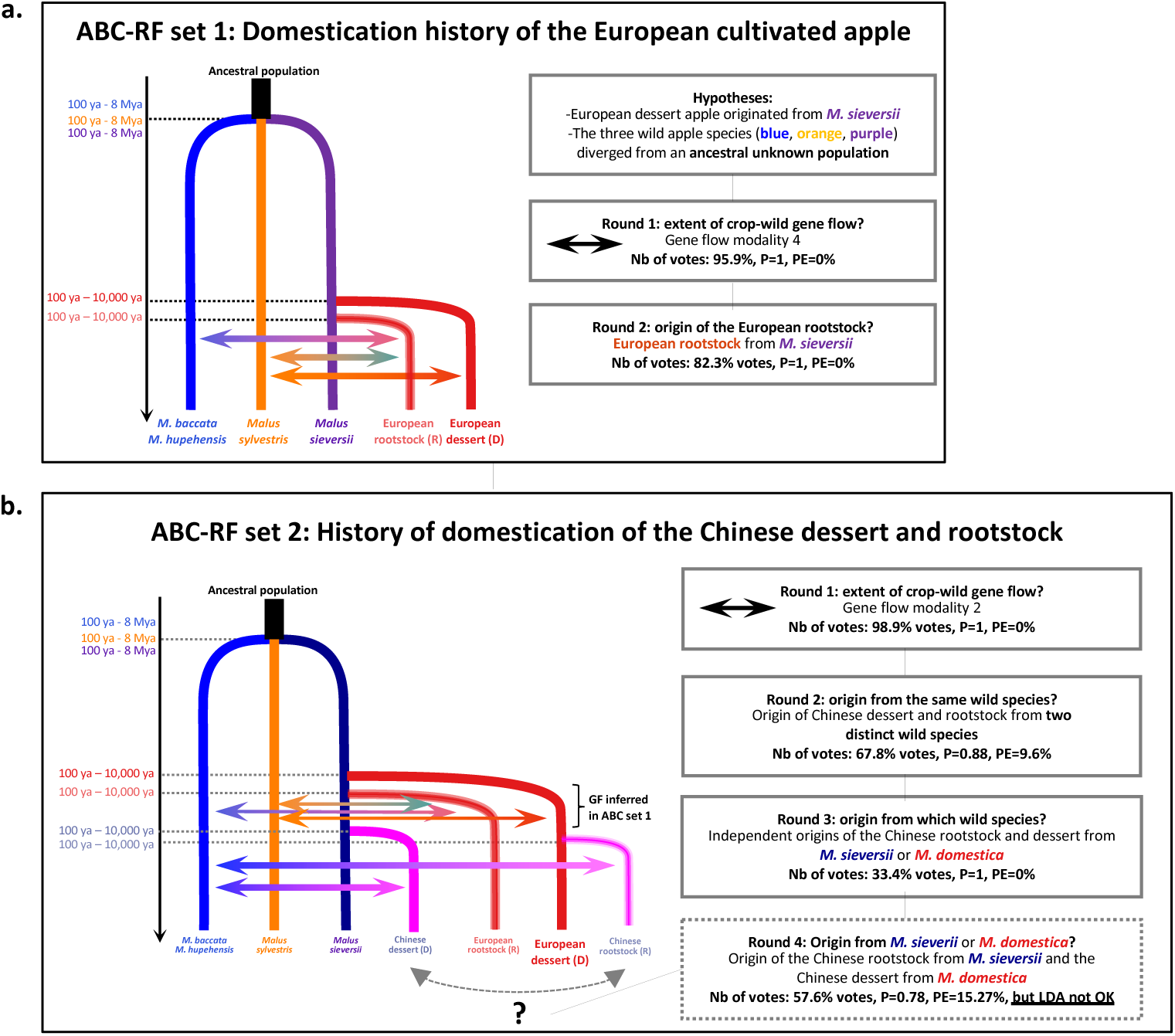
Domestication histories of European and Chinese dessert and rootstock apples inferred using the approximate Bayesian computation random forest framework (ABC-RF, 36,200 SNPs) a. Most likely scenario of domestication of the European rootstock inferred with ABC-RF set 1 (*N* = 98). b. Most likely scenario of domestication of the European and Chinese apples (set 2, *N* = 111). For each scenario, associated statistics (posterior probability, prior error rates, and percentage of votes) and parameter estimates for the final model in set 2. The grey arrow indicates that the Chinese rootstock and dessert cultivars may have originated from *M. sieversii* or *M. domestica*.

For the first round of the first ABC set (Figures 3a and S2), the classification votes were the highest 10 times out of 10 for the group of scenarios that assumed gene flow among the European dessert (*M. domestica*) and *M. sylvestris*, European rootstock and *M. sylvestris*, and *M. baccata/M. hupehensis* (*i.e.*, gene flow modality 4 in Figures S3 and 3, 479.50 votes out of the 500 RF trees, posterior probability *P* = 1.00, prior error rate = 0.00, Table S7, Figure S12a, Figure 3a). In the second round, the classification votes were the highest 10 times out of 10 for the scenarios that assumed the European rootstock from *M. sieversii* (422 votes out of the 500 RF trees, posterior probability *P* = 0.87, prior error rate = 10.02%, Table S8, Figure S12b). Thus, the first ABC set indicated the European rootstock originated from *M. sieversii* with subsequent gene flow from *M. baccata* and *M. sylvestris*.

For the first round of the second ABC set, the classification votes were the highest 10 times out of 10 for the scenarios assuming gene flow between the Chinese rootstock and dessert with *M. baccata* (*i.e.*, gene flow modality 2, 98.9% RF trees voting for this group, posterior probability *P* = 1.00, prior error rate = 0.00%, Table S9 and Figure S13a). For the second round of the second ABC set, the classification votes were the highest 10 times out of 10 for the scenarios that assumed the two Chinese cultivated apples were domesticated independently from different wild species (67.8% RF trees voting for this group, posterior probability *P* = 0.88, prior error rate = 9.61%, Table S10 and Figure S10b). For the third round, the classification votes were the highest 10 times out of 10 for the scenarios that assumed the Chinese rootstock and dessert originated from *M. sieversii* or *M. domestica* (56% RF trees voting for this group of scenarios, posterior probability *P* = 1.00, prior error rate = 0.00%, Table S11 and Figure S13c). For the fourth round, the classification votes were the highest 10 times out of 10 for the scenarios that assumed the Chinese dessert apple originated from *M. domestica* (57.6% votes out of the 500 RF trees, posterior probability *P* = 0.78, prior error rate = 15.27%, Table S12, Figure S13d). However, origin of the Chinese dessert from *M. domestica* needs to be considered with caution, as we were not able to fit the observed data within the LDA (Figure S13d).

The final scenario indicated that the European rootstock diverged from *M. sieversii* and the Chinese cultivars did not diverge from *M. baccata*, but their origin, either from *M. sieversii* or *M. domestica*, remains unclear (Figure 3b). The ABC further confirmed TreeMix gene flow estimates: *M. sylvestris* and *M. baccata* contributed to the European and Chinese dessert and rootstock gene pools (Figure 3b). The model parameters are listed in Table S13, and the posterior estimates are presented in Table S13. However, it should be noted that the credibility intervals were large and the parameter estimates need to be carefully considered. The goodness of fit of the two models (Figures S15 and S16) confirmed our model choice (*P* = 0.069 and *P* = 0.07, respectively).

## Discussion

In this study, we used population genomic approaches in combination with SNPs to investigate the relative contributions of each wild apple relatives, *M. sylvestris* and *M. sieversii*, and a supposed contributor, *M. baccata,* to the genomes of cultivated European and Chinese dessert and rootstock apples. We showed that the cultivated European dessert and rootstock apples grouped together and formed a specific gene pool, whereas the Chinese dessert and rootstock apples were a mixture of the three wild apple gene pools, mainly with *M. baccata* and *M. sieversii*. The coalescent-based inferences indicate that both European and Chinese rootstocks diverged from *M. sieversii* (or *M. domestica* for the cultivated Chinese apple gene pool), with subsequent gene flow from the wild species *M. sylvestris* and *M. baccata*. We also confirmed previous results for the contribution of *M. sylvestris* to the cultivated dessert apple gene pool (Cornille et al., 2012). Therefore, our results show that *M. baccata* is an additional contributor to the cultivated apple genome, and we have also provided insight into the origin of the apple rootstock. This study confirmed that domestication of the apple tree involved several wild apple species and that crop-wild species hybridization had a key role in fruit tree domestication. Substantial hybridization between domestic and wild forms have also been described in other fruit trees (Groppi, Liu, Cornille, Decroocq, & Decroocq, 2021; Liu et al., 2019; Wu et al., 2018), but the apple tree is the model system with the most documentation. Our results support the view that domestication of woody perennials, and crops in general, was probably a protracted and diffuse process that involved multiple geographically disparate species (Allaby, Fuller, & Brown, 2008; Purugganan, 2019).

### The European and Chinese apple rootstocks did not originate from *M. baccata*

Although grafting has been an important part of perennial woody crop evolution and breeding (Vavilov, 1926; Warschefsky et al., 2016; Zohary & Spiegel-Roy, 1975), surprisingly little is known about the origin of rootstocks. In apples, most of the studies focused on the history of dessert and cider cultivars, mostly from the Western Hemisphere. The Central Asian wild apple, *M. sieversii*, is now known to be the wild ancestor of the cultivated dessert and cider apple, *M. domestica* (Cornille et al., 2012; Cornille et al., 2019; Cornille, Giraud, Smulders, Roldán-Ruiz, & Gladieux, 2014; Daccord et al., 2017; Duan et al., 2017; Harris, Robinson, & Juniper, 2002; Migicovsky, Gardner, Richards, Thomas Chao et al., 2021; Peace et al., 2019). However, the origin of apple rootstock has not yet been elucidated. Here, we provide new insights into the domestication history of apple rootstock and show that *M. sieversii* is the progenitor of the European apple rootstock and either *M. sieversii* or *M. domestica* is the progenitor of the Chinese apple. The mixed origin of Chinese dessert cultivars, the shared ancestral polymorphism between *M. domestica* and *M. sieversii*, and the current low number of Chinese cultivars, may explain our inability to infer the origin of the Chinese dessert apple with ABC. Indeed, only four main Chinese cultivars are still cultivated in China; they are supposed to belong to two species, *M*. *domestica* subsp. *chinensis* and *M. asiatica*. Note as well that only few main rootstocks are used in China (Wang et al., 2019). In our study, the sampling size of the Chinese cultivated apple was limited because of its cultivation history in this country.

Thus, despite the large use of *M. baccata and M. hupehensis* (Wang et al., 2019) as breeding sources in China and for rootstock in Europe, *M. baccata* is not the initial progenitor of the rootstock varieties. Rather, our results suggest that *M. baccata* did contribute to apple domestication through wild-to-crop gene flow.

### Major contribution of *M. baccata* to the cultivated apple genome through crop-to-wild introgression

There is much debate on the contribution of other wild species along the Silk Route to the genetic makeup of the cultivated apple genome. Recently, the Caucasian crab apple has been shown to be an additional contributor to the cultivated apple genome (Bina et al., 2021). Our results demonstrate that interspecific hybridization has been a driving force in the evolution of apple. We confirm that the wild European crabapple *M. sylvestris* has been a major secondary contributor to the diversity of apples. We also show that the Siberian wild apple is a contributor to the diversity of the cultivated apple. Furthermore, we show the substantial gene flow from *M. baccata* to the European apple rootstock and Chinese dessert apple and rootstock. This is not surprising because *M. baccata* is widely used in the high-latitude apple-producing areas of China as a rootstock and breeding resource because of its disease resistance and cold tolerance (Chen et al., 2019; Zhi-Qin, 1999). In the Western hemisphere, *M. baccata* may be used as a parent for some crosses. Interspecific hybridization has also been observed in date palms (Flowers et al., 2019), grapes (Myles et al., 2011), almonds (Delplancke et al., 2011), and apricots (Groppi et al., 2021; Liu et al., 2019; Q. Zhang et al., 2018). Substantial crop-wild gene flow is mostly due to the self-incompatibility system of fruit trees that favors the selection of the best phenotypes grown from open-pollinated seeds and appears to be a key element in perennial crop evolution (Cornille et al., 2014; Gaut et al., 2015).

### Concluding remarks and perspectives

The domestication of fruit trees stands in stark contrast to that of annuals, especially with respect to the extent of wild-crop gene flow. Substantial crop-wild gene flow makes the resolution of relationships between crop and wild fruit trees challenging. In addition, the problem with unravelling the domestication history of fruit trees is magnified by their recent domestication in terms of number of generations and thus shared ancestral polymorphism among the crop and wild fruit trees (Cornille et al., 2014; Gaut et al., 2015; Miller & Gross, 2011). We used genomic data to distinguish between crop-wild shared ancestral polymorphism and recent gene flow and showed that the history of apples is a rare well-documented example of the evolution of a domesticated crop over a long period and involved at least four wild species. Introgression of from wild relatives is a likely source of crop adaptation (Burgarella et al., 2018). The contribution of several species to the cultivated apple genome raises questions about their role in the adaptation of cultivated apples during domestication worldwide. Genome-wide association showed that traits associated with fruit quality and texture were selected during apple domestication (Duan et al., 2017; Migicovsky, Gardner, Richards, Thomas Chao et al. 2021). However, the genomic landscape of wild-to-crop introgressions has not been investigated on the basis of genome-wide data (Daccord et al., 2017; Duan et al., 2017; Migicovsky & Myles, 2017; Sun et al., 2020; Velasco et al., 2010). Further investigations of the genomic architecture of wild-crop gene flow, and its adaptive role during domestication, are required to understand the role of gene flow during the divergence and adaptation of woody perennials to new environments.

## Acknowledgements

We thank the Franco-Chinese Campus France program « Decouverte » 2017, the *Jeunes Talents France-Chine* 2019 program, and the ATIP-Avenir CNRS Inserm grant for funding. We also thank Adrien Falce, Olivier Langella and Benoit Johannet for help and support on the INRAE-Génétique Quantitative et Evolution-Le Moulon lab cluster. We thank the INRAE MIGALE bioinformatics platform (http://migale.jouy.inra.fr) for providing help and support, in particular Véronique Martin, Eric Montaubon and Valentin Loux. This work was financially supported by the National Key Research and Development Program of China (2018YFD1000101, 2019YFD1000803); the Key Technology R & D and Industrialization Demonstration Science and Technology for the Transformation and Upgrading of Apple Industries in Shaanxi Province-Apple Anvil Combinations Evaluation and Seedlings Industrialization (2020zdzx03-01-04); The China Apple Research System (CARS 27), Tang Scholar by Cyrus Tang Foundation and Northwest A&F University.

## Data Availability

The RNAseq raw data are available at NCBI sequence read archive (SRA) under BioProject accession PRJNA763361 (https://dataview.ncbi.nlm.nih.gov/object/PRJNA763361?reviewer=5i8m9p6b4qi4ucg4kf0qe45v5j&sort_by=-accession&page=2). Vcf used for analyses is available at the Zenodo (https://zenodo.org/record/5513618). Scripts are available at https://forgemia.inra.fr/amandine.cornille/rnaseq_apple_evolutionary_history/.

## Author Contributions

DZ, AC, and XC conceived and designed the experiments; AC, DZ and MH obtained the funding; XC, NA, LX, YW sampled the material; XC performed the molecular analyses; JM and CJ performed the RNA extraction; XC and AC analyzed the data. The manuscript was written by AC and XC, with essential inputs from other co-authors.

## Notes

### Competing Interest Statement

The authors have declared no competing interest.

