## Supplementary material for "The Siberian wild apple, *Malus baccata* (L.) Borkh., is an additional contributor to the genomes of cultivated European and Chinese apples"

**Table S1.** Details for each *Malus* accession used in this study, for each filtering step.

**See Table_S1.xlsx**

Table footnotes: we referred to as cultivated group (CULT) any wild or cultivated apple species used for apple dessert or cider consumption (CULT), for ornamental (CULT_ORN) or rootstock (CULT_ROOT) purposes. Any wild species that naturally occurs in forest and are not used for human resource purposes were referred to as “WILD”. Each individual was coded accordingly: 1) WILD or CULT; 2) a 3-letter abbreviation (*e.g*., DOM for *M. domestica*, SIE for *M. sieversii*, BAC for *M. baccata*, SYL for *M. sylvestris*, ASI for *M. asiatica*, see details for other species in Tables S1 and 3) a 3-letter country or region of origin (*e.g.*, CHN for China, JPN for Japan, NZL for New-Zealand and so on, see details in supplemental dataset 1); 4) an individual ID.

**Table S2.** Sequencing data and read mapping statistics for the sequenced samples retrieved from Duan et *al*. (2017). Individual ID can be found in Table S1.

| **Individual ID** | **Raw data (G)** | **Clean data (G)** | **Raw Reads (M)** | **Clean Reads (M)** | **Mapped Reads (M)** | **Mapped rate (%)** | **Mapped Depth (X)** | **Mapped Coverage (%)** |
| --- | --- | --- | --- | --- | --- | --- | --- | --- |
| C98_Wc_China | 14.16 | 13.68 | 157.37 | 146.10 | 143.99 | 99.34% | 18.68 | 84.14% |
| C97_Wc_China | 14.77 | 14.28 | 164.15 | 154.12 | 154.11 | 99.02% | 18.61 | 80.78% |
| C99_Wc_China | 9.89 | 9.57 | 109.93 | 104.00 | 102.29 | 99.23% | 12.76 | 75.97% |
| A24_W_USA | 14.47 | 12.49 | 131.60 | 117.23 | 114.56 | 97.38% | 15.44 | 65.97% |
| A25_W_Russia | 12.53 | 10.98 | 124.04 | 109.85 | 109.08 | 98.33% | 14.44 | 81.83% |
| C108_Wc_China | 13.55 | 13.23 | 150.51 | 144.03 | 139.46 | 99.35% | 17.78 | 81.60% |
| C123_Wc_China | 18.64 | 18.18 | 207.16 | 192.28 | 193.95 | 99.05% | 24.00 | 84.53% |
| A26_W_USA | 18.32 | 14.80 | 147.37 | 118.79 | 120.36 | 98.41% | 18.79 | 78.54% |
| A10_D_Australia | 15.55 | 12.64 | 154.00 | 126.54 | 127.07 | 98.39% | 16.67 | 60.50% |
| A18_D_Canada | 22.11 | 18.53 | 176.90 | 155.54 | 153.44 | 99.04% | 25.24 | 84.44% |
| R5_D_Canada | 6.64 | 6.01 | 65.72 | 59.72 | 61.46 | 99.30% | 8.09 | 79.13% |
| R6_D_Canada | 8.71 | 7.96 | 86.19 | 80.32 | 79.28 | 99.72% | 10.90 | 81.40% |
| R2_D_CzeCh | 6.33 | 5.65 | 62.71 | 56.24 | 55.72 | 97.29% | 7.29 | 78.56% |
| A03_D_France | 25.13 | 19.86 | 190.77 | 153.81 | 155.02 | 97.73% | 25.33 | 80.55% |
| A12_D_France | 20.55 | 18.38 | 203.49 | 184.34 | 181.53 | 98.38% | 24.07 | 80.98% |
| A23_D_Germany | 16.06 | 14.88 | 160.61 | 149.43 | 145.22 | 96.66% | 19.18 | 81.72% |
| A01_D_Israel | 15.91 | 13.75 | 130.23 | 112.56 | 116.87 | 99.88% | 18.10 | 80.70% |
| A08_D_Japan | 25.21 | 22.42 | 214.57 | 190.61 | 182.48 | 92.25% | 26.44 | 80.78% |
| A09_D_NewZealand | 21.21 | 17.41 | 169.35 | 139.92 | 138.57 | 98.13% | 23.13 | 83.08% |
| R1_D_Poland | 9.35 | 7.69 | 79.49 | 70.04 | 71.61 | 98.76% | 9.99 | 75.12% |
| A02_D_Russia | 8.49 | 7.60 | 84.09 | 77.02 | 77.78 | 99.14% | 10.01 | 80.25% |
| A15_D_Russia | 25.41 | 20.09 | 207.75 | 164.69 | 168.73 | 98.78% | 25.84 | 80.03% |
| A19_D_Russia | 25.57 | 20.20 | 201.24 | 168.79 | 164.62 | 98.30% | 26.03 | 78.81% |
| A04_D_UK | 15.80 | 12.74 | 122.49 | 100.72 | 99.47 | 99.83% | 16.74 | 77.90% |
| A05_D_UK | 14.28 | 12.77 | 141.35 | 130.72 | 130.82 | 99.18% | 17.16 | 81.92% |
| A20_D_UK | 17.03 | 13.89 | 138.13 | 113.25 | 117.07 | 99.63% | 18.63 | 80.87% |
| A22_D_UK | 17.46 | 16.10 | 174.55 | 162.03 | 167.53 | 99.70% | 21.68 | 83.53% |
| R3_D_UK | 12.79 | 11.67 | 126.63 | 116.63 | 114.57 | 98.49% | 15.25 | 81.29% |
| R4_D_UK | 14.82 | 13.64 | 146.71 | 135.25 | 136.59 | 99.01% | 18.41 | 83.01% |
| R7_D_UK | 8.10 | 7.61 | 80.19 | 78.56 | 74.46 | 99.03% | 9.93 | 77.46% |
| A06_D_USA | 16.66 | 14.56 | 147.50 | 132.93 | 132.55 | 99.99% | 19.31 | 81.76% |
| A07_D_USA | 19.17 | 16.33 | 154.71 | 137.58 | 132.71 | 98.87% | 22.06 | 83.41% |
| A11_D_USA | 15.76 | 13.67 | 145.13 | 130.95 | 126.16 | 99.04% | 18.33 | 82.36% |
| A13_D_USA | 15.54 | 13.29 | 135.57 | 116.54 | 117.43 | 98.42% | 17.63 | 82.21% |
| A14_D_USA | 22.65 | 18.86 | 185.82 | 155.05 | 154.06 | 98.78% | 25.39 | 83.16% |
| A16_D_USA | 23.70 | 17.34 | 201.40 | 159.09 | 151.76 | 99.94% | 23.70 | 84.93% |
| A17_D_USA | 19.72 | 16.50 | 156.59 | 136.55 | 130.36 | 98.40% | 21.97 | 83.04% |
| A21_D_USA | 20.07 | 15.60 | 168.30 | 134.14 | 133.55 | 99.79% | 21.29 | 82.33% |
| R8_D_USA | 5.74 | 5.03 | 51.89 | 48.50 | 47.91 | 99.49% | 6.54 | 67.34% |
| A00_D_USA | 7.53 | 7.06 | 74.57 | 69.70 | 71.26 | 99.19% | 9.34 | 80.85% |
| R10_D_USA | 8.32 | 7.78 | 82.41 | 76.64 | 76.32 | 98.76% | 10.29 | 79.17% |
| R11_D_USA | 10.63 | 9.93 | 105.22 | 99.57 | 99.71 | 98.11% | 13.20 | 81.68% |
| R12_D_USA | 5.85 | 5.47 | 57.93 | 56.07 | 53.58 | 98.38% | 7.01 | 75.01% |
| A27_W_UK | 35.70 | 29.29 | 279.27 | 229.58 | 241.95 | 101.24% | 36.79 | 67.85% |
| A29_W_USA | 13.36 | 9.79 | 132.23 | 99.76 | 95.99 | 97.33% | 12.16 | 33.58% |
| C113_Wc_China | 15.66 | 15.25 | 174.04 | 166.57 | 166.56 | 99.27% | 19.97 | 79.04% |
| A30_W_USA | 11.41 | 9.15 | 112.93 | 92.84 | 87.05 | 95.55% | 11.15 | 63.75% |
| C106_Wc_China | 13.28 | 12.88 | 147.51 | 137.37 | 140.62 | 99.15% | 16.67 | 80.36% |
| C102_Wc_China | 12.75 | 12.38 | 141.70 | 131.01 | 131.17 | 99.37% | 16.61 | 81.88% |
| A32_W_Turkey | 20.18 | 16.39 | 167.08 | 142.85 | 137.49 | 99.00% | 22.19 | 83.88% |
| C104_Wc_China | 13.96 | 13.61 | 155.15 | 145.26 | 148.03 | 99.28% | 18.10 | 81.78% |
| A33_W_USA | 20.46 | 16.91 | 167.36 | 145.40 | 137.56 | 98.83% | 22.11 | 80.12% |
| A34_W_Kazakhstan | 18.03 | 14.56 | 167.12 | 138.14 | 132.06 | 97.30% | 18.87 | 76.59% |
| C105_Wc_China | 15.28 | 14.83 | 169.75 | 161.24 | 159.06 | 98.90% | 19.56 | 83.78% |
| C80_S_Xinjiang | 15.69 | 15.29 | 174.35 | 168.33 | 163.12 | 98.57% | 20.17 | 80.72% |
| C86_S_Xinjiang | 16.30 | 15.92 | 181.06 | 171.19 | 170.05 | 99.49% | 21.52 | 81.89% |
| C87_S_Xinjiang | 16.17 | 15.78 | 179.67 | 167.16 | 162.67 | 97.40% | 21.09 | 82.66% |
| C78_S_Xinjiang | 13.98 | 13.74 | 155.35 | 149.67 | 144.60 | 99.55% | 18.17 | 79.85% |
| C81_S_Xinjiang | 13.81 | 13.58 | 153.50 | 145.40 | 144.87 | 98.53% | 18.05 | 81.13% |
| C88_S_Xinjiang | 14.06 | 13.82 | 156.21 | 151.00 | 149.07 | 99.26% | 18.70 | 82.01% |
| C90_S_Xinjiang | 13.55 | 13.30 | 150.57 | 144.58 | 145.44 | 99.30% | 17.39 | 79.13% |
| C74_S_Xinjiang | 14.60 | 14.31 | 162.26 | 155.57 | 151.37 | 99.34% | 18.76 | 79.39% |
| C77_S_Xinjiang | 12.22 | 11.99 | 135.83 | 129.50 | 130.87 | 99.48% | 16.30 | 81.93% |
| C85_S_Xinjiang | 16.03 | 15.70 | 178.07 | 172.66 | 164.79 | 99.37% | 21.00 | 80.98% |
| C89_S_Xinjiang | 16.72 | 16.40 | 185.75 | 180.43 | 172.99 | 98.61% | 22.28 | 83.06% |
| C93_S_Xinjiang | 15.32 | 14.67 | 170.19 | 161.82 | 161.21 | 99.10% | 19.59 | 82.18% |
| C94_S_Xinjiang | 18.93 | 18.47 | 210.37 | 204.70 | 202.85 | 99.40% | 24.58 | 81.26% |
| C96_S_Xinjiang | 13.93 | 13.53 | 154.82 | 147.39 | 142.88 | 99.18% | 18.56 | 84.18% |
| A35_S_Kazakhstan | 17.93 | 13.81 | 145.36 | 118.31 | 116.69 | 99.14% | 18.45 | 80.97% |
| A36_S_Kazakhstan | 15.39 | 13.44 | 131.66 | 115.76 | 114.94 | 98.71% | 17.49 | 79.67% |
| A37_S_Kazakhstan | 16.02 | 12.99 | 135.57 | 114.20 | 109.24 | 97.89% | 17.16 | 81.98% |
| A38_S_Kazakhstan | 17.70 | 14.99 | 150.18 | 132.66 | 126.44 | 98.69% | 19.53 | 80.38% |
| A39_S_Kazakhstan | 16.66 | 13.88 | 141.08 | 122.15 | 119.26 | 98.33% | 18.28 | 78.72% |
| A40_S_Kazakhstan | 14.53 | 12.03 | 125.24 | 105.64 | 107.44 | 98.93% | 16.08 | 81.37% |
| A41_S_Kazakhstan | 16.79 | 14.59 | 143.19 | 128.25 | 124.84 | 98.87% | 18.98 | 80.38% |
| A42_S_Kazakhstan | 16.05 | 14.02 | 138.30 | 121.18 | 124.68 | 98.34% | 18.83 | 83.56% |
| A43_S_Kazakhstan | 15.52 | 13.41 | 133.95 | 121.66 | 118.61 | 98.75% | 17.24 | 78.69% |
| A44_S_Kazakhstan | 16.78 | 13.39 | 137.47 | 114.40 | 115.21 | 99.25% | 17.93 | 80.46% |
| A45_S_Kazakhstan | 15.93 | 12.95 | 129.84 | 110.88 | 107.72 | 98.93% | 16.94 | 79.49% |
| A46_S_Kazakhstan | 12.45 | 11.19 | 104.45 | 93.79 | 94.02 | 99.34% | 14.76 | 81.02% |
| A47_S_Kazakhstan | 15.93 | 12.79 | 129.30 | 108.87 | 105.45 | 99.31% | 17.24 | 81.75% |
| A48_S_Kazakhstan | 15.60 | 13.54 | 134.74 | 121.09 | 117.58 | 99.05% | 18.26 | 83.45% |
| A49_S_Kazakhstan | 15.68 | 13.34 | 130.48 | 113.60 | 114.78 | 99.01% | 17.95 | 82.93% |
| A31_W_Russia | 21.04 | 17.25 | 167.62 | 140.20 | 142.02 | 99.59% | 22.67 | 79.37% |
| C101_Wc_China | 14.29 | 13.91 | 158.83 | 147.12 | 149.07 | 98.07% | 18.48 | 83.54% |
| A51_St_Germany | 16.65 | 12.87 | 134.85 | 109.16 | 108.33 | 99.12% | 16.78 | 78.71% |
| A52_St_Germany | 11.73 | 10.38 | 104.94 | 95.95 | 93.90 | 98.41% | 13.60 | 75.16% |
| A53_St_Germany | 15.45 | 13.14 | 128.53 | 110.72 | 111.96 | 98.62% | 17.78 | 82.70% |
| A54_St_Germany | 14.64 | 12.71 | 125.61 | 112.04 | 112.08 | 99.13% | 16.97 | 81.17% |
| A58_St_Norway | 11.19 | 10.22 | 110.83 | 103.98 | 101.65 | 97.97% | 13.30 | 80.92% |
| A59_St_Russia | 19.31 | 15.27 | 150.80 | 120.97 | 125.31 | 99.53% | 20.43 | 82.76% |
| A56_St_UK | 18.42 | 15.08 | 151.49 | 126.67 | 128.43 | 99.44% | 20.14 | 81.63% |
| A50_St_USA | 15.82 | 13.46 | 133.80 | 116.08 | 114.01 | 99.11% | 18.33 | 83.96% |
| A55_St_Yugoslavia | 15.33 | 13.05 | 129.75 | 114.73 | 113.19 | 99.33% | 17.45 | 81.30% |
| A57_St_Yugoslavia | 19.05 | 14.82 | 156.02 | 127.98 | 126.28 | 97.99% | 19.88 | 84.26% |
| C107_Wc_China | 15.18 | 14.82 | 168.62 | 158.92 | 160.60 | 99.43% | 19.62 | 83.00% |
| C100_Wc_China | 14.96 | 14.49 | 166.26 | 155.99 | 150.78 | 96.65% | 18.44 | 81.24% |

**Table S3.** Sequencing statistics and read mapping statistics for new 71 samples sequenced for RNA added in this study. Individual ID can be found in Table S1.

| **Individual ID** | **Raw data (G)** | **Clean data (G)** | **Raw Reads (M)** | **Clean Reads (M)** | **Mapped Reads (M)** | **Mapped rate (%)** | **Depth(X)** | **Coverage (%)** |
| --- | --- | --- | --- | --- | --- | --- | --- | --- |
| D-46 | 7.58 | 6.88 | 45.24 | 41.04 | 76.71 | 93.46% | 11.00 | 14.27% |
| D-47 | 7.98 | 7.24 | 47.62 | 43.18 | 80.95 | 93.73% | 11.70 | 13.21% |
| D-48 | 6.91 | 6.25 | 41.23 | 37.30 | 68.83 | 92.27% | 10.03 | 14.89% |
| D-49 | 6.44 | 5.82 | 38.43 | 34.74 | 62.20 | 89.51% | 9.82 | 13.00% |
| D-50 | 5.25 | 4.78 | 31.30 | 28.50 | 54.01 | 94.75% | 7.47 | 13.06% |
| D-52 | 7.35 | 6.58 | 43.87 | 39.26 | 72.72 | 92.62% | 10.56 | 13.64% |
| D-53 | 5.75 | 5.19 | 34.31 | 30.98 | 57.17 | 92.26% | 8.44 | 12.41% |
| D-54 | 6.82 | 6.15 | 40.69 | 36.71 | 68.75 | 93.65% | 10.10 | 14.22% |
| D-55 | 5.00 | 4.55 | 29.84 | 27.15 | 51.93 | 95.61% | 7.02 | 11.79% |
| D-56 | 6.77 | 6.12 | 40.41 | 36.49 | 69.16 | 94.75% | 9.66 | 13.75% |
| D-57 | 6.55 | 5.85 | 39.10 | 34.90 | 65.11 | 93.28% | 9.39 | 12.24% |
| D-58 | 6.75 | 6.11 | 40.28 | 36.44 | 68.88 | 94.52% | 9.38 | 13.01% |
| D-59 | 6.10 | 5.50 | 36.38 | 32.80 | 65.01 | 99.10% | 8.42 | 11.57% |
| D-60 | 7.37 | 6.67 | 43.98 | 39.81 | 74.80 | 93.96% | 10.55 | 12.96% |
| D-62 | 7.56 | 6.83 | 45.07 | 40.75 | 72.84 | 89.38% | 11.40 | 14.86% |
| D-63 | 5.83 | 5.26 | 34.75 | 31.38 | 57.08 | 90.94% | 8.82 | 12.73% |
| D-64 | 6.30 | 5.68 | 37.59 | 33.89 | 61.79 | 91.17% | 9.50 | 13.27% |
| D-65 | 7.81 | 7.02 | 46.59 | 41.90 | 76.39 | 91.17% | 11.81 | 14.51% |
| D-66 | 6.25 | 5.51 | 37.28 | 32.89 | 59.81 | 90.92% | 9.17 | 13.02% |
| R-67 | 5.98 | 5.27 | 35.69 | 31.46 | 60.03 | 95.40% | 8.80 | 10.73% |
| R-68 | 6.63 | 5.95 | 39.56 | 35.49 | 66.42 | 93.59% | 8.86 | 12.03% |
| R-69 | 6.85 | 6.15 | 40.88 | 36.66 | 67.76 | 92.41% | 9.92 | 14.59% |
| R-70 | 6.95 | 6.24 | 41.49 | 37.21 | 68.12 | 91.54% | 10.29 | 14.50% |
| R-71 | 6.37 | 5.70 | 37.97 | 34.03 | 61.97 | 91.05% | 9.26 | 12.72% |
| R-72 | 5.60 | 5.06 | 33.41 | 30.17 | 54.00 | 89.48% | 7.90 | 14.25% |
| R-75 | 6.48 | 5.82 | 38.64 | 34.73 | 62.84 | 90.48% | 9.42 | 14.57% |
| R-77 | 7.60 | 6.70 | 45.34 | 39.96 | 72.39 | 90.56% | 11.06 | 14.20% |
| W-01 | 5.47 | 5.06 | 32.65 | 30.16 | 53.95 | 89.43% | 8.30 | 12.40% |
| W-02 | 6.10 | 5.62 | 36.38 | 33.54 | 63.10 | 94.06% | 8.75 | 13.91% |
| W-03 | 6.22 | 5.76 | 37.09 | 34.35 | 62.96 | 91.63% | 9.40 | 13.10% |
| W-04 | 6.50 | 6.03 | 38.80 | 35.96 | 66.01 | 91.79% | 9.87 | 14.09% |
| W-05 | 5.30 | 4.89 | 31.63 | 29.20 | 55.52 | 95.09% | 8.01 | 13.05% |
| W-06 | 5.35 | 4.94 | 31.91 | 29.46 | 55.42 | 94.07% | 8.06 | 13.89% |
| W-07 | 4.42 | 4.09 | 26.38 | 24.40 | 44.44 | 91.06% | 6.74 | 11.26% |
| W-08 | 5.63 | 5.18 | 33.57 | 30.89 | 56.28 | 91.10% | 8.44 | 12.74% |
| W-09 | 4.77 | 4.40 | 28.46 | 26.24 | 48.50 | 92.41% | 7.24 | 13.45% |
| W-10 | 5.26 | 4.85 | 31.38 | 28.95 | 52.54 | 90.76% | 7.94 | 14.15% |
| W-11 | 5.26 | 4.86 | 31.37 | 29.02 | 54.12 | 93.25% | 7.92 | 12.36% |
| W-12 | 6.60 | 6.19 | 39.40 | 36.93 | 71.30 | 96.53% | 9.73 | 13.08% |
| W-13 | 5.71 | 5.33 | 34.08 | 31.82 | 62.97 | 98.96% | 8.04 | 10.97% |
| W-14 | 7.34 | 6.87 | 43.76 | 40.97 | 78.68 | 96.02% | 10.49 | 13.40% |
| W-15 | 5.50 | 5.16 | 32.83 | 30.79 | 58.16 | 94.45% | 7.91 | 13.17% |
| W-16 | 7.16 | 6.71 | 42.73 | 40.02 | 76.84 | 96.01% | 10.49 | 13.58% |
| W-17 | 6.49 | 6.07 | 38.70 | 36.19 | 71.39 | 98.63% | 8.93 | 10.54% |
| W-18 | 5.85 | 5.44 | 34.87 | 32.42 | 62.96 | 97.08% | 8.13 | 10.33% |
| W-19 | 6.21 | 5.80 | 37.05 | 34.62 | 66.39 | 95.87% | 8.67 | 12.75% |
| W-20 | 4.61 | 4.43 | 27.51 | 26.42 | 50.76 | 96.06% | 6.99 | 12.37% |
| W-21 | 6.59 | 6.10 | 39.32 | 36.40 | 62.61 | 86.00% | 8.77 | 10.70% |
| W-22 | 6.09 | 5.63 | 36.30 | 33.60 | 65.43 | 97.37% | 8.46 | 12.18% |
| W-23 | 5.83 | 5.41 | 34.78 | 32.28 | 57.95 | 89.75% | 9.00 | 14.19% |
| W-24 | 4.88 | 4.62 | 29.08 | 27.59 | 54.01 | 97.89% | 6.87 | 10.99% |
| W-25 | 4.72 | 4.52 | 28.16 | 26.98 | 49.90 | 92.48% | 6.95 | 12.22% |
| W-26 | 4.98 | 4.77 | 29.68 | 28.46 | 56.69 | 99.61% | 7.04 | 10.91% |
| W-27 | 5.90 | 5.45 | 35.18 | 32.51 | 61.14 | 94.05% | 8.88 | 13.51% |
| W-28 | 5.02 | 4.64 | 29.97 | 27.66 | 54.88 | 99.18% | 7.32 | 12.48% |
| W-29 | 5.91 | 5.22 | 35.26 | 31.16 | 58.40 | 93.69% | 8.11 | 13.96% |
| W-30 | 6.52 | 5.79 | 38.90 | 34.52 | 64.36 | 93.21% | 8.68 | 12.75% |
| W-31 | 5.30 | 4.66 | 31.62 | 27.78 | 54.39 | 97.91% | 7.34 | 11.64% |
| W-32 | 5.48 | 4.84 | 32.71 | 28.86 | 53.85 | 93.31% | 7.70 | 12.56% |
| W-33 | 7.58 | 6.67 | 45.21 | 39.80 | 73.65 | 92.53% | 10.54 | 15.21% |
| W-34L | 7.18 | 6.63 | 42.81 | 39.53 | 70.95 | 89.73% | 11.00 | 15.03% |
| W-35 | 6.93 | 6.10 | 41.36 | 36.37 | 67.35 | 92.59% | 9.82 | 13.23% |
| W-36 | 6.39 | 5.54 | 38.13 | 33.07 | 62.24 | 94.09% | 9.04 | 12.92% |
| W-37L | 6.11 | 5.58 | 36.43 | 33.29 | 59.86 | 89.90% | 7.89 | 14.07% |
| W-39 | 7.09 | 6.24 | 42.27 | 37.25 | 70.77 | 95.00% | 9.48 | 12.23% |
| W-40 | 6.93 | 6.10 | 41.33 | 36.37 | 66.66 | 91.63% | 9.95 | 12.40% |
| W-41L | 6.16 | 5.71 | 36.73 | 34.05 | 62.28 | 91.45% | 8.29 | 12.26% |
| W-42L | 5.84 | 5.41 | 34.84 | 32.27 | 60.80 | 94.20% | 8.06 | 13.11% |
| W-43L | 6.36 | 5.90 | 37.96 | 35.20 | 50.45 | 71.66% | 7.13 | 9.93% |
| W-44 | 8.07 | 7.30 | 48.12 | 43.56 | 74.63 | 85.67% | 10.57 | 11.90% |
| W-45 | 6.84 | 6.16 | 40.80 | 36.77 | 71.74 | 97.56% | 9.21 | 12.16% |

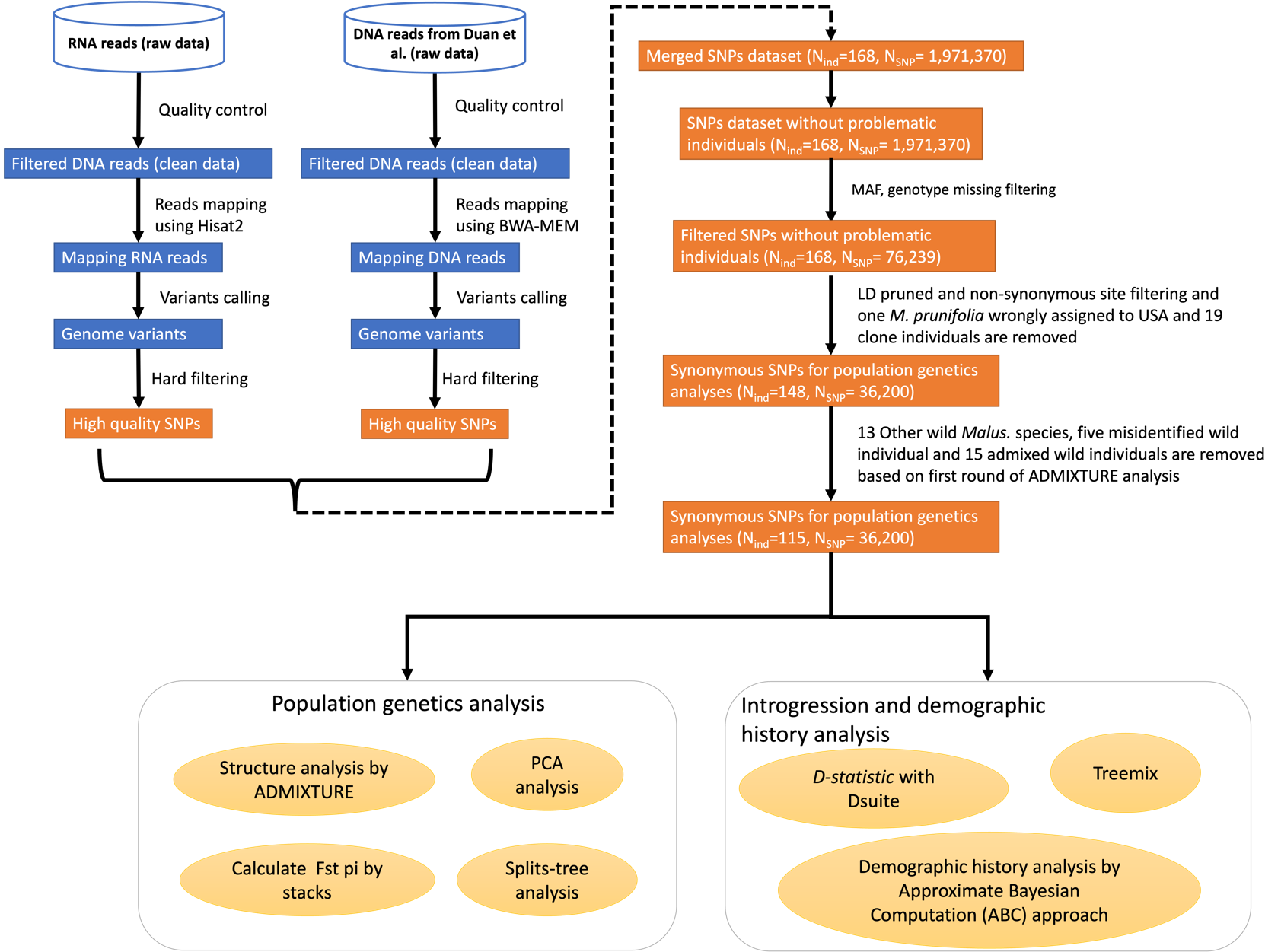

**Figure S1**. Bioinformatic pipeline for SNP calling and filtering, population genomics analyses, gene flow estimates, demographic history inferences used to reconstruct the domestication history of the European and Chinese cultivated apples.

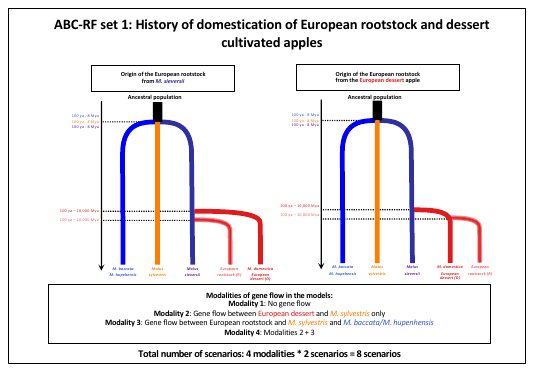

**Figure S2. Domestication history of European apple dessert and rootstock inferred using the approximate Bayesian computation random forest framework (ABC-RF set 1, *N*=98, 36,200 SNPs).** Two scenarios of domestication of the European apple rootstock tested in the ABC-RF set 1. We did not test the origin of the European dessert, as its history is already known (Cornille *et al.*, 2012, 2014; Duan *et al.*, 2017; Migicovsky *et al.*, 2021)*.* In all models, the three wild apple populations (*M. sieversii*, *M. sylvestris* and *M. baccata*-*M. hupenhensis*) diverged from a common unknown ancestral population. For the two scenarios, we assumed four modalities of gene flow. In round 1, we compared eight scenarios were compared to infer the modalities of crop-wild gene flow. We took the best model chosen in round 1, and then we compared the two divergence histories of European apple rootstock in round 2.

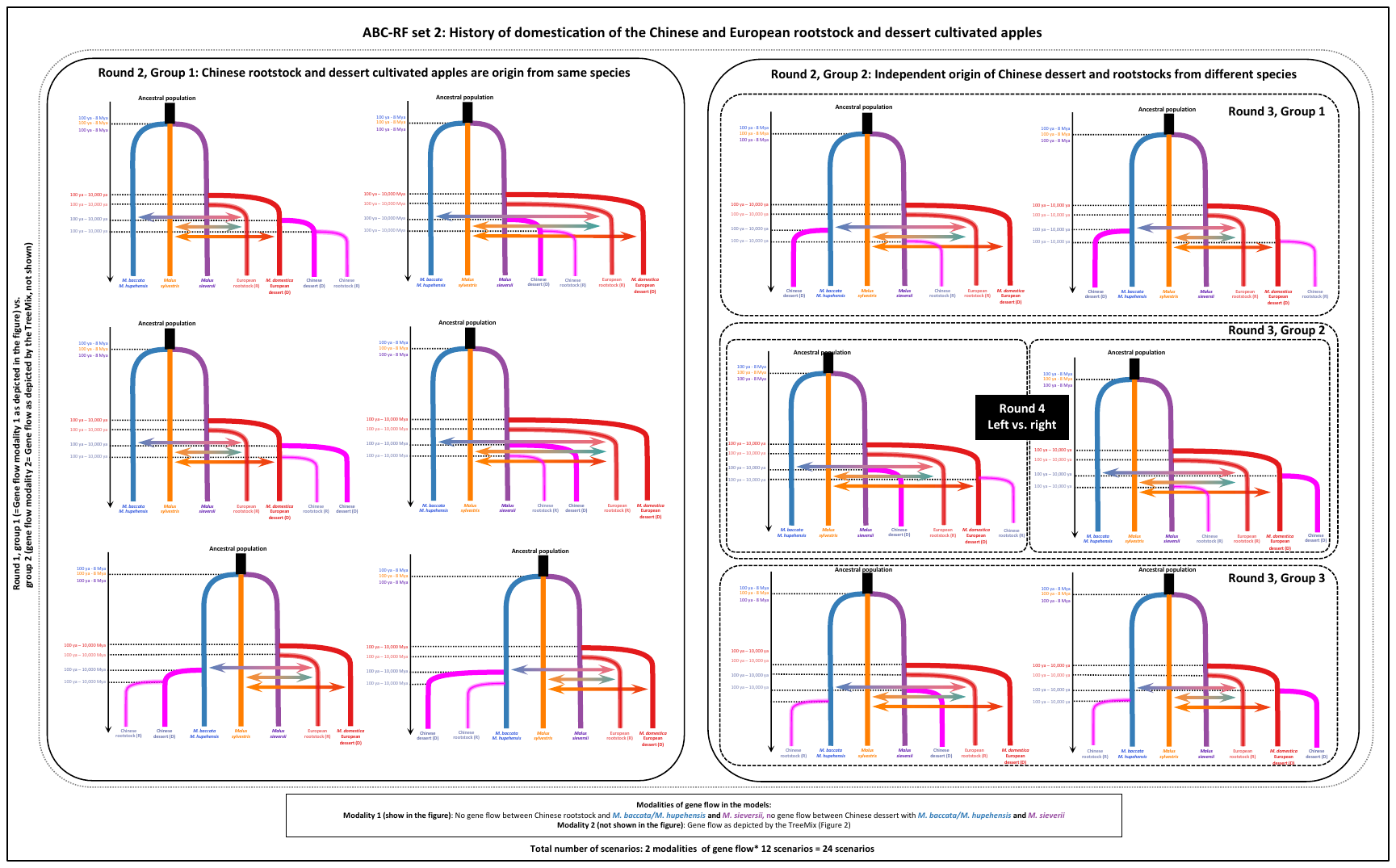

**Figure S3.** **Domestication history of the Chinese dessert and rootstock apples inferred using the approximate Bayesian computation random forest framework (ABC-RF set 2, *N*=111, 36,200 SNPs).** Once the best model in ABC-RF set 1 inferred, we added the Chinese apple cultivars. Similar to the previous ABC-RF set 1, we assumed that the three wild apples diverged from an unknown ancestral population. In the round 1, we compared 24 scenarios to test two modalities of crop-wild gene flow. In round 2, we compared 12 scenarios to tested whether the Chinese dessert and rootstock apple diverged independently from two distinct wild populations. In round 3, we tested whether the Chinese rootstock and dessert apples diverged from the *M. baccata-M. hupehensis* population. In round 4, we compared the two scenarios as depicted in the figure (right vs. left) which tested the origin of the Chinese dessert apple from M*. sieversii* or *M. domestica*.

**Table S4.** Prior distributions used for approximate Bayesian computations for inferring the domestication history of the European and Chinese cultivated apples.

|  | Parameter | Distribution | Lower bound | Upper bound |
| --- | --- | --- | --- | --- |
| ABC analyses set1 and set2 | *N_X_** | uniform | 50 | 50,000 |
|  | *N_ANC_* | *N_SYL_ + N_BAC_HUP_ + N_SIE_* | | |
|  | *T_X-ANC_* | log uniform | 100 | 8,000,000 |
|  | *T_X-Y_* | log uniform | 100 | 8,000,000 |
|  | *T_C-X_* | log uniform | 100 | 10,000 |
|  | *M_X-Y_* | uniform | 0 | 0.5 |

Note: Prior distributions are uniform and log uniform between lower and upper bound. *N_X_*: effective population size of population *X*; *T_X-ANC_*: Divergence time between population *X* and an unknown ancestral population ANC; Divergence time were calculated by multiplying the generation time estimates of *Malus* species (5-12 years (Crosby *et al.*, 1990) we assumed a generation time of 10 years.); *M_X-Y_*: migration rates from populations *X* to *Y*; *T_X-Y_*: Divergence time between populations *X* and *Y*. *T_X-C_* : Divergence time between wild populations *X* and cultivated populations. * *Nx* were log-transformed for fastsimcoal2 simulations.

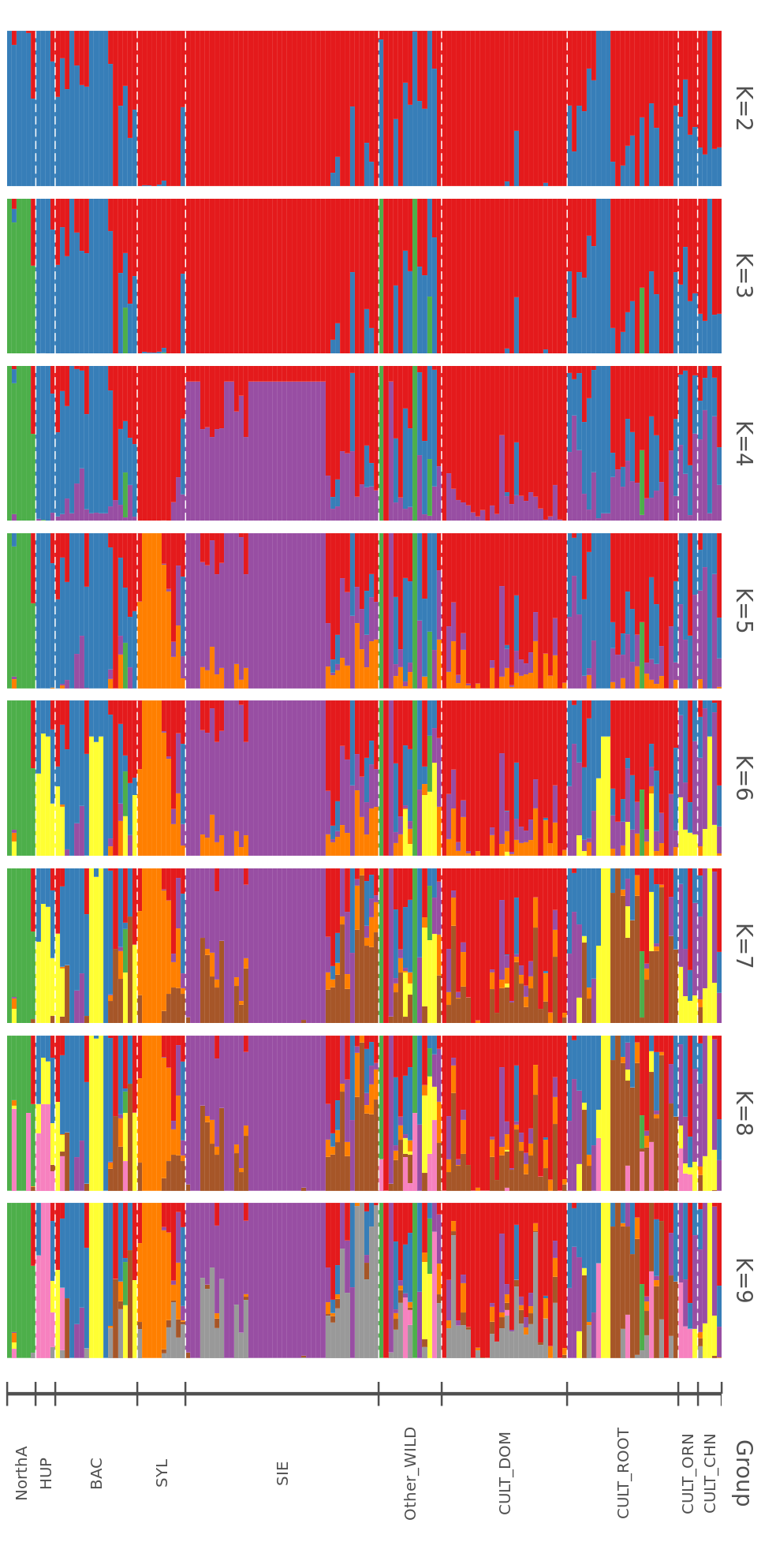

**Figure S4.** **Population genetic structure among the wild and cultivated apples inferred from *K*=2 to *K*=9 with ADMIXTURE (*N*=148, 36,200 SNPs).** NorthA: wild apple individuals from North America (*N*=6, *i.e.*, *Malus angustifolia, Malus coronaria, Malus fusca, Malus ioensis,* and *Malus transitoria*); HUP: *Malus hupehensis* (*N*=4); BAC: *Malus baccata* (*N*=17); SYL: *Malus sylvestris* (*N*=10), SIE: *Malus sieversii* (*N*=40), Other WILD (*N*=13, *i.e.*, *Malus florentina, Malus halliana, Malus honanensis, Malus kansuensis, Malus micromalus, Malus orientalis, Malus sikkimensis, Malus soulardii, Malus toringoides, Malus x platycarpa,* and *Malus yunanensis*); CULT_DOM: *Malus domestica* (*N*=26), CULT_ROOT: Apple cultivars used as rootstocks (*N*=23, *i.e., Malus domestica, Malus prunifolia,* and *Malus robusta*); CULT_ORN: apple cultivars used for ornamental purpose (*N*=4), *i.e.*, *Malus floribunda, Malus micromalus, Malus spectabilis,* and *Malus zumi*); CULT_CHN: apple cultivars from China (*N*=5), *i.e.,* *Malus asiatica* and hybrid species).

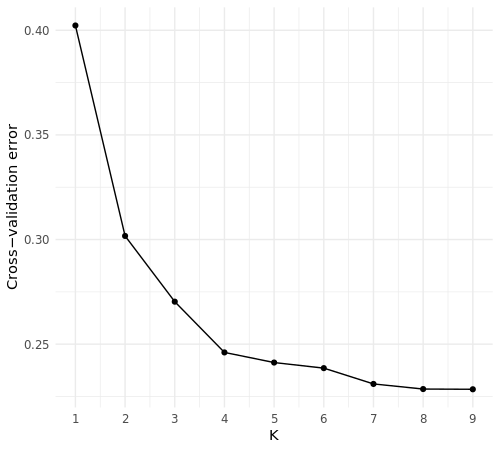

**Figure S5.** Cross-validation plot for each *K* values obtained with ADMIXTURE analyses for the wild and cultivated apples (*N*=148, 36,200 SNPs).

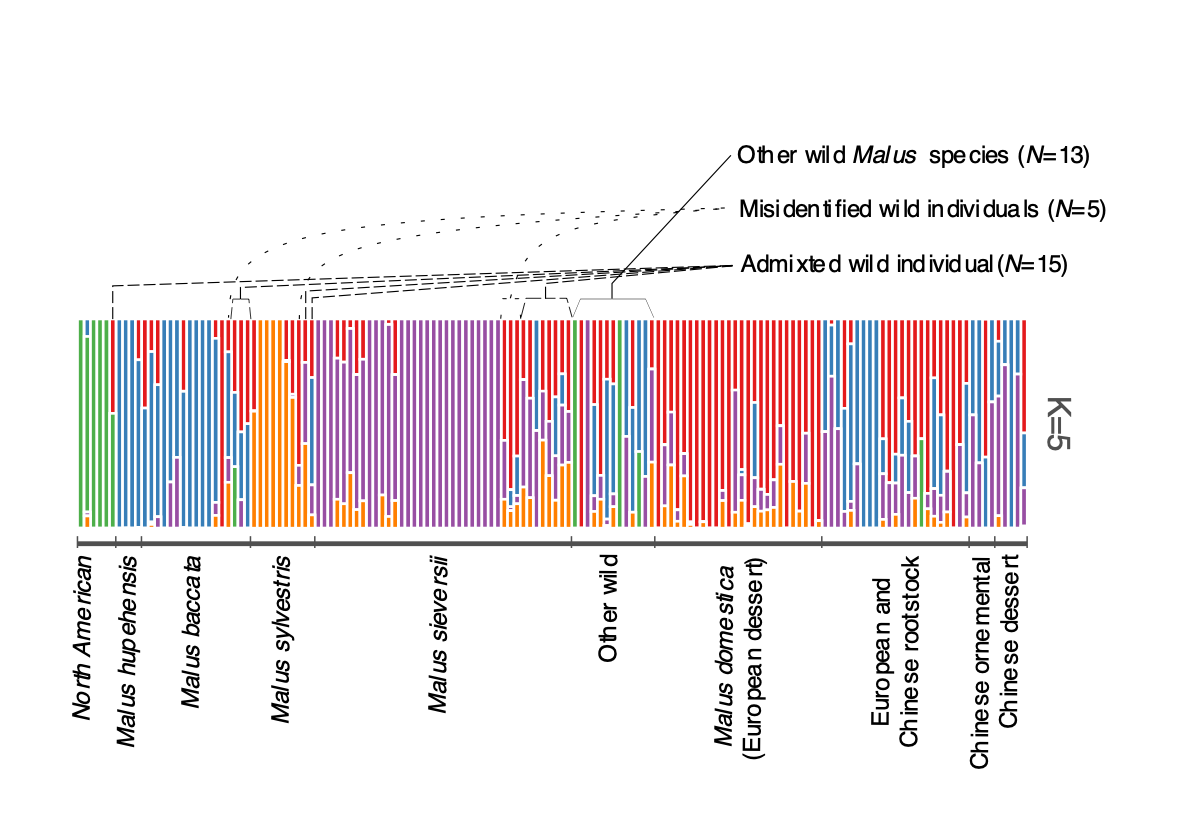

**Figure S6.** Population structure for *K*=5 for the wild and cultivated apples for the whole dataset (*i.e.*, after the removal of clones and one individual with an unknown origin: *N*=148, 36,200 SNPs), and explanation of the filtering of individuals to obtain the final pruned dataset (after removing the wild admixed or misidentified individuals, *N*=115, 36,200 SNPs). The group names are as in Table 1.

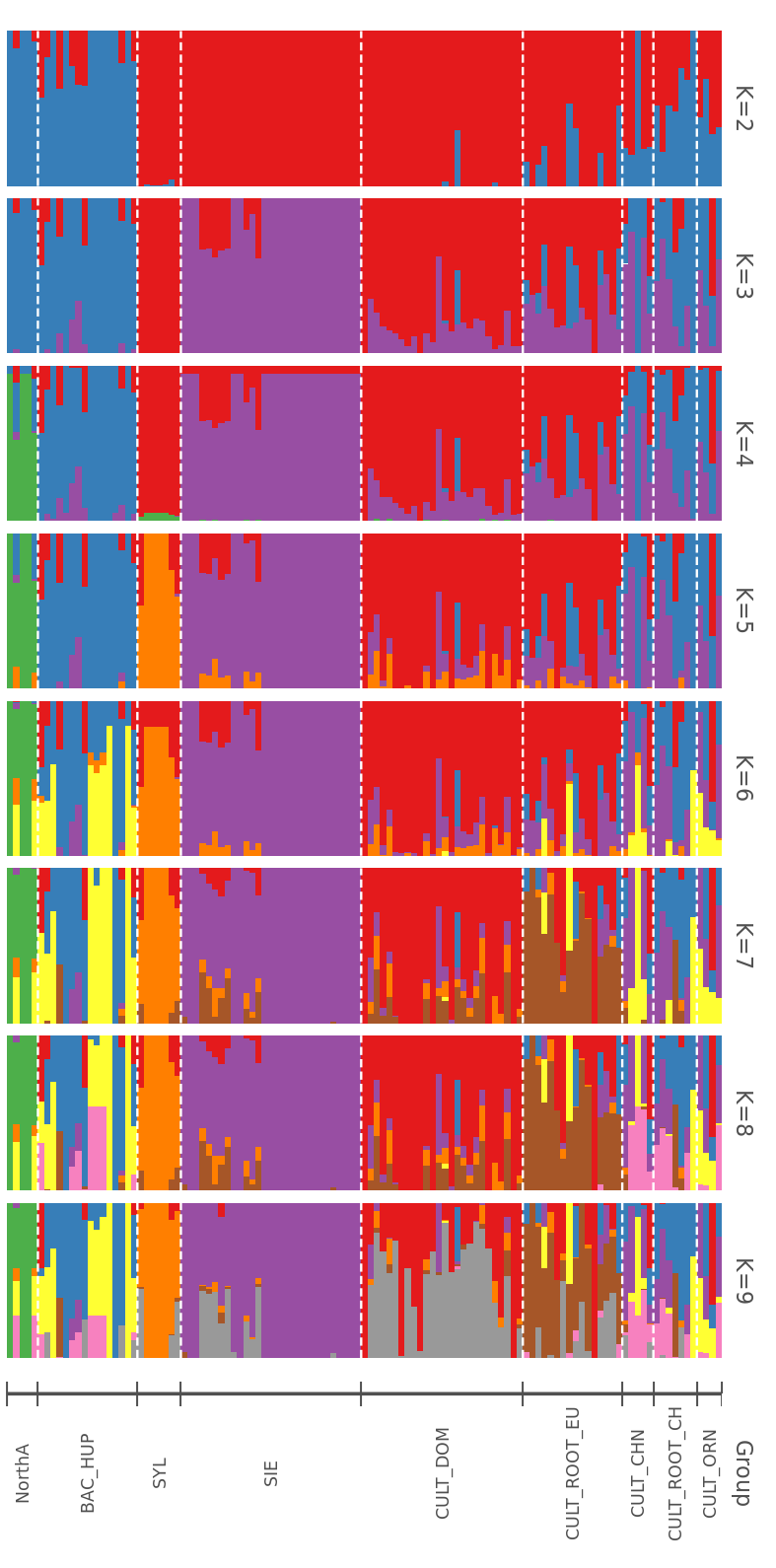

**Figure S7. Population genetic structure among the wild and the cultivated apples inferred from *K*=2 to *K*=9 with ADMIXTURE (final pruned dataset, *N*=115, 36,200 SNPs).** NorthA: *Malus* individuals from North America (*N*=6, *i.e.*, *Malus angustifolia, Malus coronaria, Malus fusca, Malus ioensis,* and *Malus transitoria*), HUP_BAC: *Malus hupehensis* and *Malus baccata* (*N*=20), SYL: *Malus sylvestris* (*N*=7), SIE: *Malus sieversii* (*N*=29), CULT_DOM: *Malus domestica* (European dessert cultivars, *N*=26), CULT_ROOT_CH: Chinese rootstock apple cultivars (*N*=7, *i.e.*, *Malus prunifolia,* and *Malus robusta*); CULT_ROOT_EU: European rootstock cultivars (*N*=16), CULT_ORN: Chinese ornamental apple cultivars (*N*=4, *i.e.*, *Malus floribunda, Malus micromalus, Malus spectabilis,* and *Malus zumi*), CULT_CHN: Chinese dessert apple cultivated apple (*N*=5, *i.e.*, *Malus asiatica* and hybrid species).

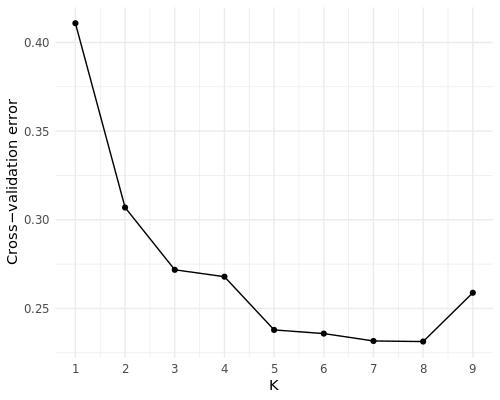

**Figure S8.** Cross-validation plot for each *K* for ADMIXTURE analysis for the wild and cultivated apples (final pruned dataset, *N*=148, 36,200 SNPs).

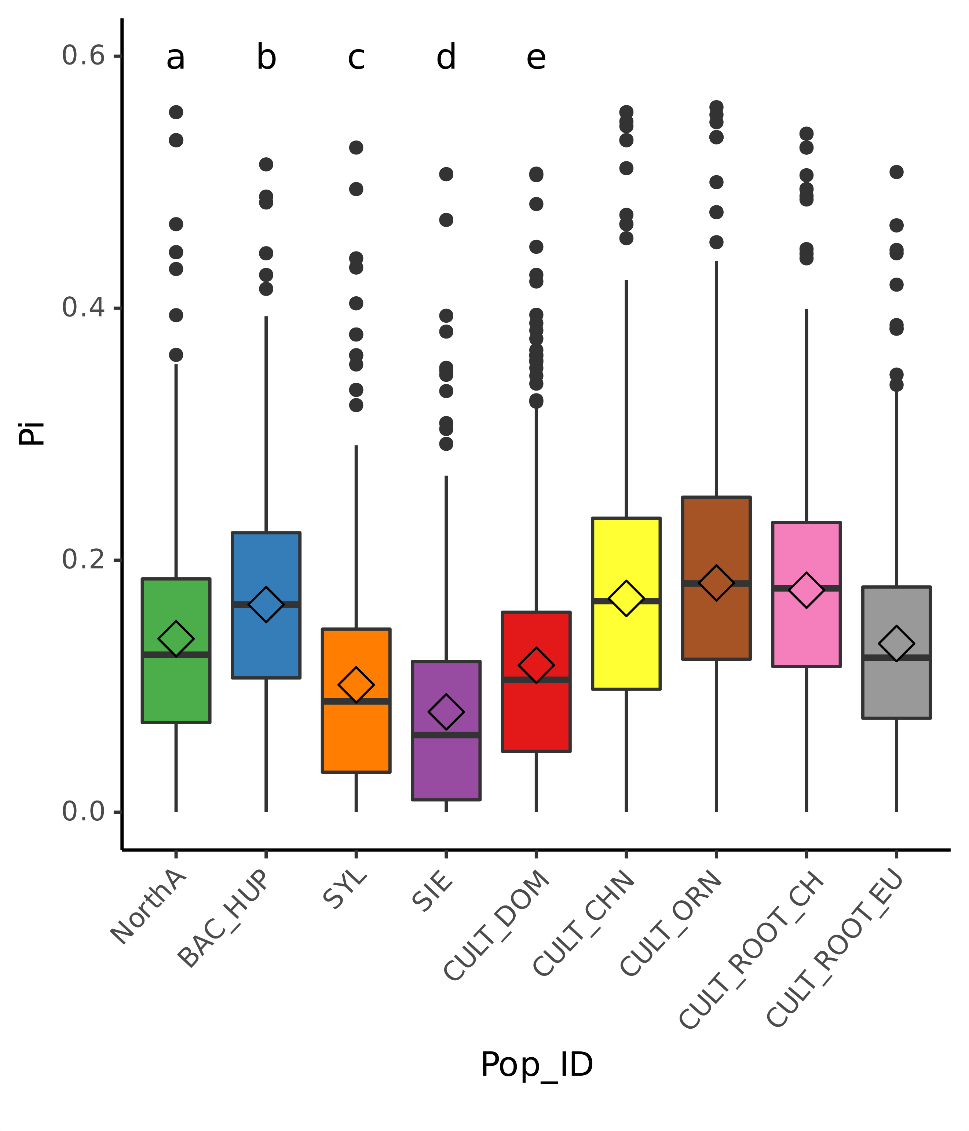

**Figure S9**. **Nucleotide diversity (Nei’s *П*) based on 50-kb non-overlapping window for the nine populations** **(*N*=115,** **36,200 SNPs**)**.** In the box plots, a black line within the box indicates the median, a diamond within the box indicates the mean, small letters show significant differences (*P-value* < 0.05 by ANOVA). Colors represent populations described in Table 1 and Figure 1. NorthA: *Malus* individuals from North America (*N*=5, *i.e.*, *Malus angustifolia, Malus coronaria, Malus fusca, Malus ioensis,* and *Malus transitoria*), HUP_BAC: *Malus hupehensis* and *Malus baccata* (*N*=16), SYL: *Malus sylvestris* (*N*=7), SIE: *Malus sieversii* (*N*=29), CULT_DOM: *Malus domestica* (European dessert cultivars, *N*=26), CULT_ROOT_CH: Chinese rootstock apple cultivars (*N*=7, *i.e.*, *Malus prunifolia,* and *Malus robusta*); CULT_ROOT_EU: European rootstock cultivars (*N*=16), CULT_ORN: Chinese ornamental apple cultivars (*N*=4, *i.e.*, *Malus floribunda, Malus micromalus, Malus spectabilis,* and *Malus zumi*), CULT_CHN: Chinese dessert apple cultivated apple (*N*=5, *i.e.*, *Malus asiatica* and hybrid species).

**Table S5.** Genetic diversity estimates for the wild and cultivated apple populations inferred at *K*=5 with ADMIXTURE (*N*=115, 36,200 SNPs).

| Status | Population | N | *H_O_* | *H_E_* | *F_IS_* | *π* |
| --- | --- | --- | --- | --- | --- | --- |
| Wild | North America | 5 | 0.0541 | 0.1174 | 0.1598** | 0.1304** |
|  | *M. baccata/M. hupehensis* | 16 | 0.1564 | 0.1561 | 0.0160** | 0.1611** |
|  | *Malus sylvestris* | 7 | 0.0846 | 0.0848 | 0.0179** | 0.0913** |
|  | *Malus sieversii* | 29 | 0.0742 | 0.0728 | 0.0066** | 0.0741** |
|  | Average (wild) | 57 | 0.09 | 0.11 | 0.05 | 0.11 |
| Crop | European dessert | 26 | 0.1136 | 0.1092 | 0.0005** | 0.1113** |
|  | Chinese dessert | 5 | 0.1584 | 0.1531 | 0.0307** | 0.1701** |
|  | Chinese ornamental | 4 | 0.2172 | 0.1617 | -0.0611** | 0.1848** |
|  | Chinese rootstock | 7 | 0.2051 | 0.166 | -0.0555** | 0.1788** |
|  | European rootstock | 16 | 0.1414 | 0.1308 | -0.0111** | 0.1350** |
|  | Average (crop) | 58 | 0.18 | 0.15 | -0.02 | 0.17 |

Note: *H_O_* and *H_E_*: observed and expected heterozygosities, respectively, *F_IS_*: inbreeding coefficient, π: Nei’s nucleotide diversity. **: *P < 0.001*.

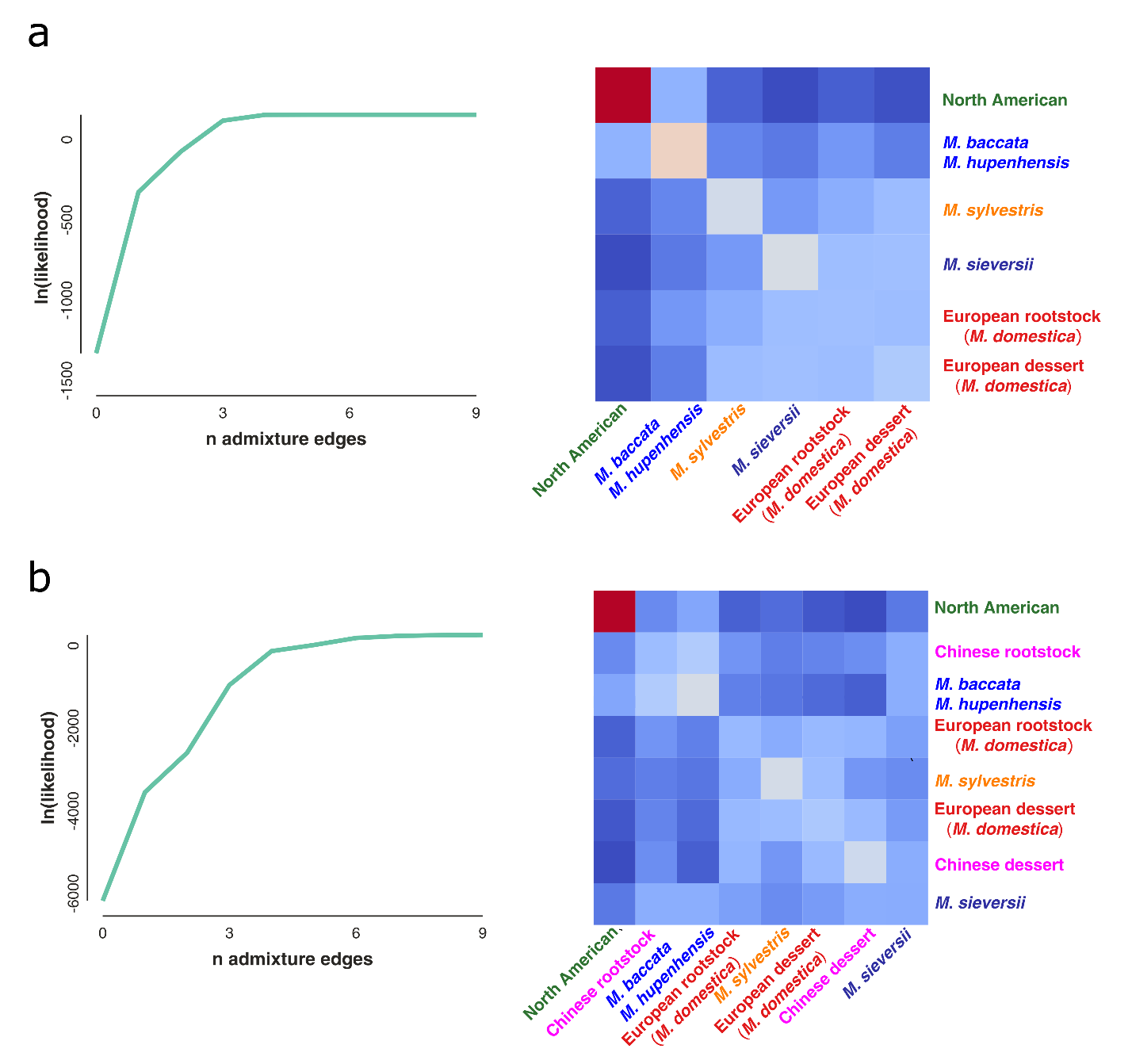

**Figure S10.** Log likelihood score of each model fit (assuming 0 to 9 migration events) and covariance matrix among eight wild and cultivated apple populations (the Chinese ornamental, *N*=4, were removed because of the small number of individuals, Table 1) inferred with TreeMix.

**Table S6**. Patterson’s *D* statistic (ABBA-BABA statistic) estimated with D-suite used to assess evidence of gene flow among the wild apple species and Chinese and European cultivated apples.

| P1 | P2 | P3 | *C_ABBA_* | *C_BABA_* | *C_BBAA_* | *D-statistic* | *Z-score* | *P-value* |
| --- | --- | --- | --- | --- | --- | --- | --- | --- |
| **SIE** | **CULT_DOM** | **SYL** | **1148.92** | **883.84** | **477.79** | **0.30** | **13.29** | **0.00** |
| **CULT_ROOT_EU** |  | **SYL** | **1004.44** | **946.56** | **606.20** | **0.22** | **17.72** | **0.00** |
| SYL |  | BAC_HUP | 2089.37 | 487.34 | 402.92 | 0.09 | 9.59 | 0.00 |
| **SYL** | **CULT_ROOT_EU** | **BAC_HUP** | **1805.58** | **680.60** | **459.48** | **0.19** | **16.73** | **0.00** |
| **SIE** |  | **BAC_HUP** | **2146.91** | **575.33** | **434.42** | **0.14** | **11.06** | **0.00** |
| **CULT_DOM** |  | **BAC_HUP** | **2117.30** | **594.07** | **457.37** | **0.13** | **11.53** | **0.00** |
| **SYL** | **CULT_CHN** | **BAC_HUP** | **1277.02** | **1063.69** | **550.00** | **0.32** | **27.79** | **0.00** |
| **SIE** |  | **BAC_HUP** | **1839.41** | **918.55** | **485.08** | **0.31** | **21.92** | **0.00** |
| **CULT_DOM** |  | **BAC_HUP** | **1581.38** | **983.56** | **554.29** | **0.28** | **24.89** | **0.00** |
| CULT_ROOT_CH |  | SYL | 1119.66 | 1024.69 | 698.24 | 0.19 | 9.82 | 0.00 |
| **CULT_ROOT_EU** |  | **BAC_HUP** | **1504.69** | **949.04** | **656.47** | **0.18** | **15.54** | **0.00** |
| **SIE** | **CULT_ROOT_CH** | **BAC_HUP** | **1373.76** | **1290.22** | **577.44** | **0.38** | **26.66** | **0.00** |
| BAC_HUP |  | SYL | 1385.59 | 993.18 | 592.60 | 0.25 | 17.70 | 0.00 |
| **CULT_CHN** |  | **BAC_HUP** | **1134.44** | **1105.44** | **826.14** | **0.14** | **9.62** | **0.00** |

Note: P1, P2, P3: three populations used in the test, details of the populations are described in Table 1; *C_ABBA_, C_BABA_ and C_BBAA_*: count of the ABBA, BABA and BBAA sites. *Z-score* and *P-value* associated with the *D-statistics* are computed over 20 jack-knife blocks divided the dataset; The bolded rows are consistent with the results of TreeMix. Population ID are described in Figure S9.

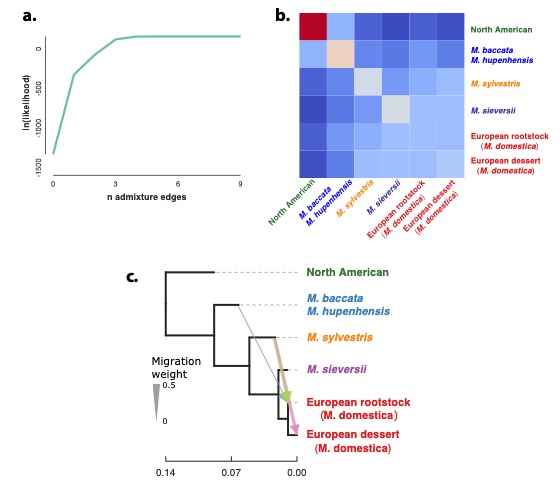

**Figure S11. TreeMix analysis ran for settling one of the gene flow modalities of the ABC set 1 to infer the history of the European apple rootstock.** a. Log likelihood score of each model fit (assuming 0 to 9 migration events) and covariance matrix among four wild and the two European cultivated apple populations (the Chinese cultivars were removed, Table 1); the best number of migration events was three. c. European rootstock underwent gene flow from *M. baccata/M. hupenhensis* and *M. sylvestris*, European dessert *M. domestica* cultivars underwent gene flow from *M. sylvestris*.

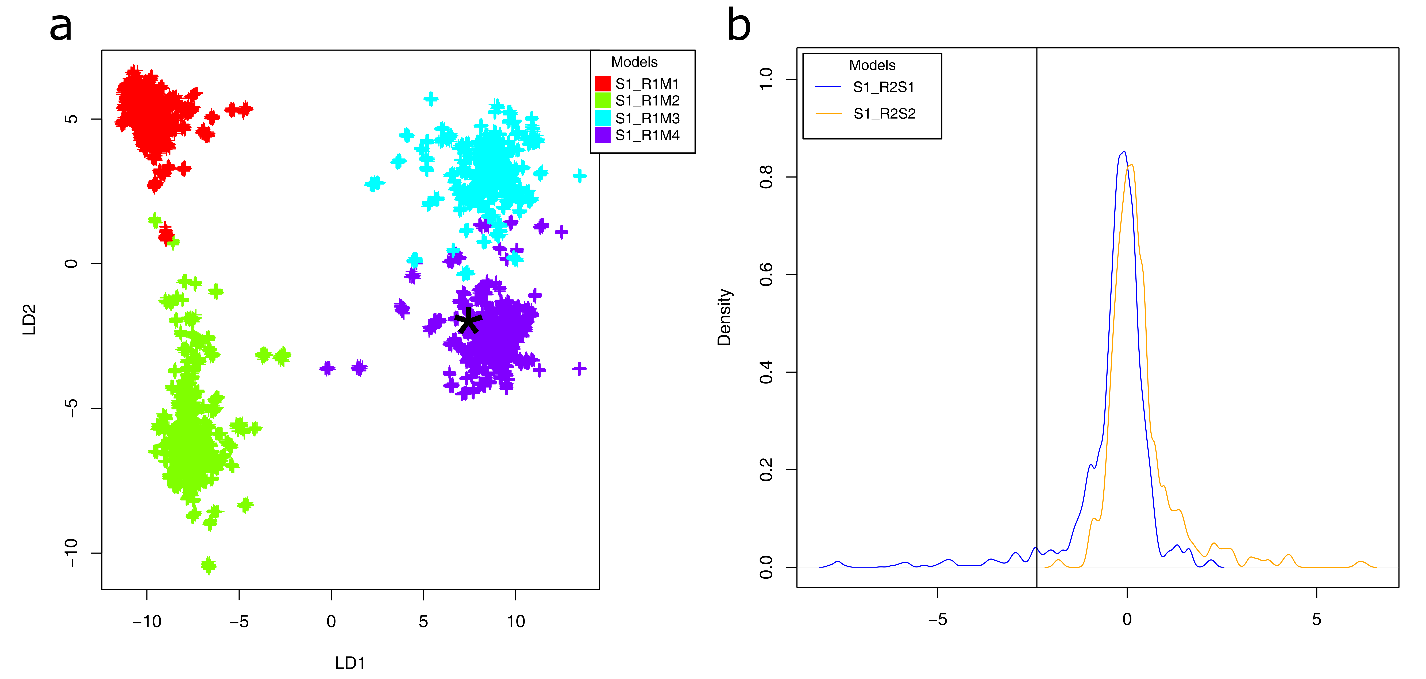

Figure S12. **Visual assessment of Random-Forest approximate Bayesian computation analyses comparing scenarios of domestication history of the European *Malus domestica* rootstocks (ABC, set 1).** a. Projection of the simulated summary statistics obtained from the reference table on a single Linear Discriminant analysis (LDA) axes for round 1. The first round included eight scenarios with two scenarios of divergence the European rootstock cultivars, simulated with i) no gene flow (S1_R1M1), ii) bidirectional gene flow between *M. domestica* and *M. sylvestris* (S1_R1M2), iii) bidirectional gene flow between the European rootstock and *M. sylvestris*, and European rootstock and *M. baccata/M. hupehensis* (S1_R1M3), iv) bidirectional gene flow between *M. domestica* and *M. sylvestris*, and bidirectional gene flow between the European rootstock and *M. sylvestris*, and European rootstock and *M. baccata/M. hupehensis* (S1_R1M4). b. Projection of the simulated summary statistics obtained from the reference table on a single LDA axes for round 2. The second round tested the divergence history of the European rootstock depicted in Figure 3a, S1_R2S1: the European rootstock diverged from *M. sieversii*, S1_R2S1: the European rootstock diverged from *M. domestica* cultivated apples. The black asterisk or black vertical line represents the observed data, and each cross represents one simulation.

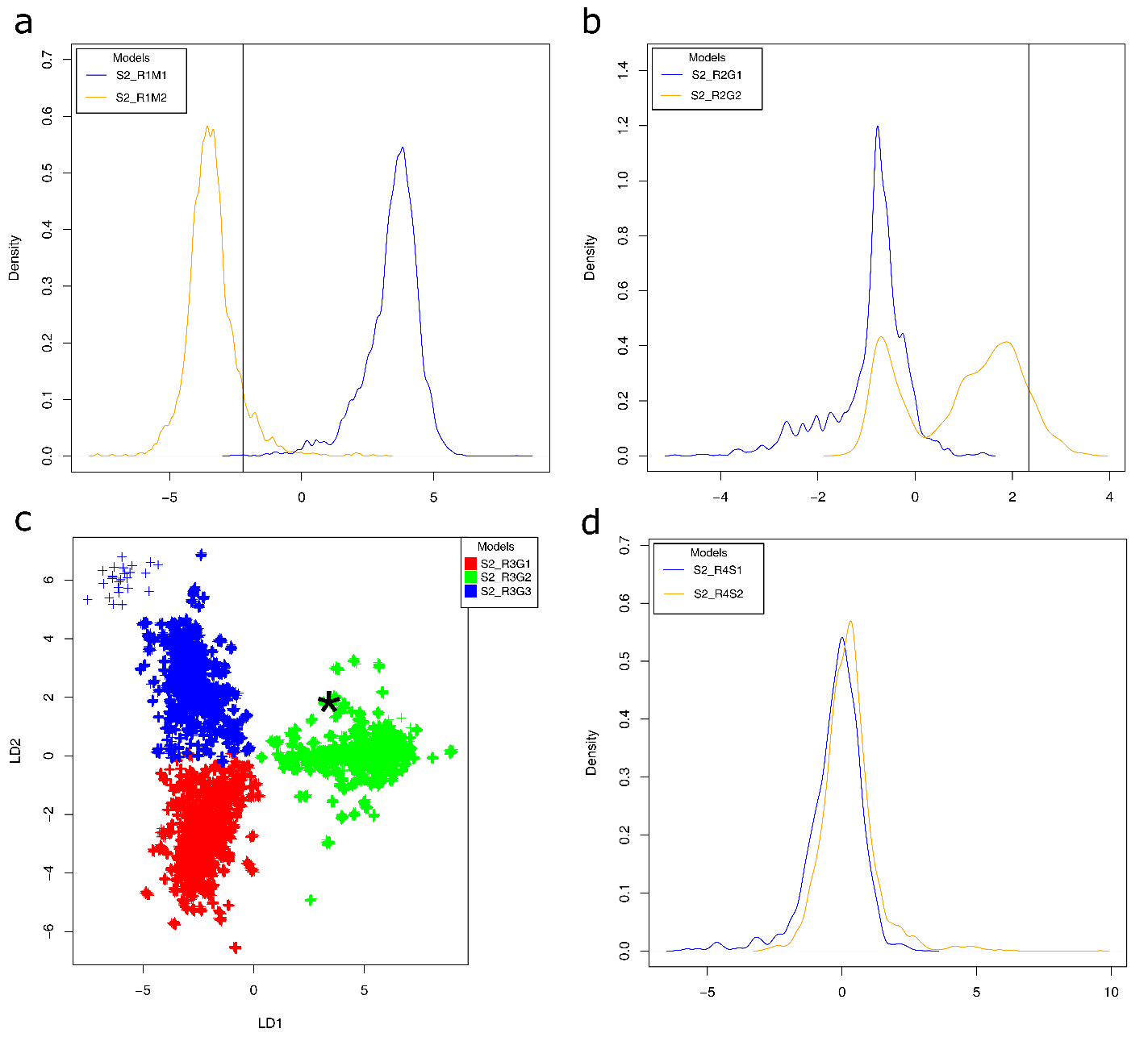

Figure S13 **Visual assessment of Random-Forest approximate Bayesian computation analyses comparing scenarios of domestication history of European and Chinese dessert and rootstock apples (ABC, set 2).** a. Projection of the simulated summary statistics obtained from the reference table in a Linear Discriminant analysis (LDA) for round 1. The first round included 24 scenarios including 12 scenarios divergence of the Chinese dessert and rootstock cultivated apples, simulated i) without gene flow among Chinese rootstock and dessert apple with *M. baccata/M. hupehensis* and *M. sieversii* (S2_R1M1), ii) with bidirectional gene flow between the Chinese rootstock and dessert apple and the wild apples as depicted by TreeMix (Figure 2, S2_R1M1). b. Projection of the simulated summary statistics obtained from the reference table on a single Linear Discriminant analysis axes for round 2. i) Chinese rootstock and dessert cultivated apples originated from the same species. ii) Independent origins of Chinese dessert and rootstocks from different species. (c) Projection of the simulated summary statistics obtained from the reference table on a single LDA axes for round 3. i) Independent origins of Chinese dessert from *M. baccata/M. hupehensis,* and an origin of Chinese rootstock from *M. sieversii* or *M. domestica*; ii) Independent origins of Chinese rootstock and dessert from *M. sieverii* and *M. domestica; iii)* Independent origins of Chinese rootstock from *M. baccata/M. hupehensis,* and origin of Chinese dessert from *M. sieversii* or *M. domestica*. Each scenario was simulated 8,000 times. The black asterisk or black vertical line represents the observed data, and each cross represents one simulation.

**Table S7.** Results of the ABC-RF algorithm to infer the European dessert and rootstock demographic histories (set 1, round 1, Figures 3a) assuming four groups of scenarios described in Figure 3a and Figure S12a. The most likely model is **S1_R1M4** scenario group (ten out of ten).

| Replicate | Votes number of different scenarios | | | | Posterior probability | Prior error rate (%) |
| --- | --- | --- | --- | --- | --- | --- |
|  | S1_R1M1 | S1_R1M2 | S1_R1M3 | **S1_R1M4** |  |  |
| 1 | 0 | 1 | 21 | **478** | 1 | 0 |
| 2 | 0 | 2 | 15 | **483** | 1 | 0 |
| 3 | 0 | 1 | 20 | **479** | 1 | 0 |
| 4 | 0 | 1 | 21 | **478** | 1 | 0 |
| 5 | 0 | 2 | 25 | **473** | 1 | 0 |
| 6 | 1 | 3 | 14 | **482** | 1 | 0 |
| 7 | 0 | 2 | 17 | **481** | 1 | 0 |
| 8 | 0 | 0 | 15 | **485** | 1 | 0 |
| 9 | 0 | 1 | 23 | **476** | 1 | 0 |
| 10 | 0 | 2 | 18 | **480** | 1 | 0 |
| Mean | 0.1 | 1.5 | 18.9 | **479.5** | 1 | 0 |
| Standard deviation | 0.32 | 0.85 | 3.7 | **3.5** | 0 | 0 |

Note: Posterior probability and prior error rate is for the best scenario with the greatest number of votes.

**Table S8.** Results of the ABC-RF algorithm comparing the cultivated apple rootstocks evolutionary histories (set 1 round 2, Figures 3a) assuming two scenarios described in Figures S2 and S12b. The most likely model is **S1_R2S1** scenario (ten out of ten).

| Replicate | Votes number of different scenarios | | Posterior probability | Prior error rate (%) |
| --- | --- | --- | --- | --- |
|  | **S1_R2S1** | S1_R2S2 |  |  |
| 1 | **416** | 84 | 0.89 | 10.13 |
| 2 | **408** | 92 | 0.83 | 10.26 |
| 3 | **425** | 75 | 0.88 | 10 |
| 4 | **423** | 77 | 0.88 | 9.83 |
| 5 | **429** | 71 | 0.87 | 10.08 |
| 6 | **418** | 82 | 0.88 | 10.12 |
| 7 | **422** | 78 | 0.86 | 9.85 |
| 8 | **420** | 80 | 0.87 | 9.85 |
| 9 | **437** | 63 | 0.86 | 10.08 |
| 10 | **422** | 78 | 0.89 | 10.04 |
| **Mean** | **422** | 78 | 0.87 | 10.02 |
| **Standard deviation** | **7.72** | 7.72 | 0.02 | 0.14 |

Note: Posterior probability and prior error rate are for the best scenario with the greatest number of votes.

**Table S9.** Results of the ABC-RF algorithm comparing the history of domestication of the Chinese rootstock and dessert apples (set 2, round 1) assuming two groups of scenarios with four different gene flow modalities described in Figures 3b and Figure S13a. The most likely model is **S2_R1M2** scenario group (ten out of ten).

| Replicate | Votes number of different scenarios | | Posterior probability | Prior error rate (%) |
| --- | --- | --- | --- | --- |
|  | S2_R1M1 | **S2_R1M2** |  |  |
| 1 | 4 | **496** | 1.00 | 0.00 |
| 2 | 0 | **500** | 1.00 | 0.00 |
| 3 | 10 | **490** | 1.00 | 0.00 |
| 4 | 5 | **495** | 1.00 | 0.00 |
| 5 | 5 | **495** | 1.00 | 0.00 |
| 6 | 1 | **499** | 1.00 | 0.00 |
| 7 | 13 | **487** | 1.00 | 0.00 |
| 8 | 4 | **496** | 1.00 | 0.00 |
| 9 | 6 | **494** | 1.00 | 0.00 |
| 10 | 7 | **493** | 1.00 | 0.00 |
| **Mean** | 5.50 | **494.50** | 1.00 | 0.00 |
| **Standard deviation** | 3.87 | **3.87** | 0.00 | 0.00 |

Note: Posterior probability and prior error rate are for the best scenario with the greatest number of votes.

**Table S10.** Results of the ABC-RF algorithm comparing history of domestication of the Chinese rootstock and dessert cultivated apples (set 2, round 2) described in Figure S3b and S13b. The most likely model is **S2_R2G2** scenario group (ten out of ten).

| Replicate | Votes number of different scenarios | | Posterior probability | Prior error rate (%) |
| --- | --- | --- | --- | --- |
|  | S2_R2G1 | **S2_R2G2** |  |  |
| 1 | 167 | **333** | 0.87 | 9.62 |
| 2 | 148 | **352** | 0.88 | 9.68 |
| 3 | 170 | **330** | 0.89 | 9.62 |
| 4 | 152 | **348** | 0.88 | 9.61 |
| 5 | 177 | **323** | 0.88 | 9.61 |
| 6 | 168 | **332** | 0.88 | 9.58 |
| 7 | 161 | **339** | 0.87 | 9.57 |
| 8 | 150 | **350** | 0.89 | 9.61 |
| 9 | 158 | **342** | 0.92 | 9.62 |
| 10 | 161 | **339** | 0.88 | 9.61 |
| **Mean** | 161.2 | **338.8** | 0.88 | 9.61 |
| **Standard deviation** | 9.44 | **9.44** | 0.01 | 0.03 |

Note: Posterior probability and prior error rate is for the best scenario with the greatest number of votes.

**Table S11.** Results of the ABC-RF algorithm comparing history of domestication of the Chinese rootstock and dessert apples (set 2, round 3) described in Figures 3b and S13c. The most likely model is **S2_R3G2** scenario group (ten out of ten).

| Replicate | Votes number of different scenarios | | | Posterior probability | Prior error rate (%) |
| --- | --- | --- | --- | --- | --- |
|  | S2_R3G1 | **S2_R3G2** | S2_R3G1 |  |  |
| 1 | 94 | **279** | 127 | 1.00 | 0.00 |
| 2 | 89 | **287** | 124 | 1.00 | 0.00 |
| 3 | 108 | **265** | 127 | 1.00 | 0.00 |
| 4 | 108 | **280** | 112 | 1.00 | 0.00 |
| 5 | 106 | **278** | 116 | 1.00 | 0.00 |
| 6 | 93 | **289** | 118 | 1.00 | 0.00 |
| 7 | 93 | **278** | 129 | 1.00 | 0.00 |
| 8 | 95 | **286** | 119 | 1.00 | 0.00 |
| 9 | 85 | **280** | 135 | 1.00 | 0.00 |
| 10 | 118 | **278** | 104 | 1.00 | 0.00 |
| Mean | 98.9 | **280** | 121.1 | 1.00 | 0.00 |
| Standard deviation | 10.44 | **6.70** | 9.12 | 0.00 | 0.00 |

Note: Posterior probability and prior error rate is for the best scenario with the greatest number of votes.

**Table S12.** Results of the ABC-RF algorithm comparing history of domestication of the Chinese rootstock and dessert apples (set 2, round 4) described in Figures 3b and S13d. The most likely model is **S2_R4S2** scenario (ten out of ten).

| Replicate | Votes number of different scenarios | | Posterior probability | Prior error rate (%) |
| --- | --- | --- | --- | --- |
|  | S2_R4S1 | **S2_R4S2** |  |  |
| 1 | 217 | **283** | 0.79 | 15.51 |
| 2 | 217 | **283** | 0.75 | 15.18 |
| 3 | 215 | **285** | 0.80 | 15.13 |
| 4 | 210 | **290** | 0.80 | 15.15 |
| 5 | 189 | **311** | 0.78 | 15.20 |
| 6 | 215 | **285** | 0.78 | 15.42 |
| 7 | 209 | **291** | 0.76 | 15.48 |
| 8 | 228 | **272** | 0.75 | 15.20 |
| 9 | 198 | **302** | 0.81 | 15.13 |
| 10 | 220 | **280** | 0.74 | 15.33 |
| Mean | 211.8 | **288.2** | 0.78 | 15.27 |
| Standard deviation | 11.18 | **11.18** | 0.02 | 0.15 |

Note: Posterior probability and prior error rate is for the best scenario with the greatest number of votes.

**Table S13**. Parameters estimates inferred for the two most likely scenarios of the domestication history of the European and Chinese cultivated apples.

| Parameter | | Mean posteriori probability | | | Mean posteriori probability in years | | | q5% | | | q5% in years | | | q95% | | | q95% in yesrs | | | NMAE | | |
| --- | --- | --- | --- | --- | --- | --- | --- | --- | --- | --- | --- | --- | --- | --- | --- | --- | --- | --- | --- | --- | --- | --- |
| S2_R4S1 | S2_R4S2 | S2_R4S1 | S2_R4S2 | Averge | S2_R4S1 | S2_R4S2 | Averge | S2_R4S1 | S2_R4S2 | Averge | S2_R4S1 | S2_R4S2 | Averge | S2_R4S1 | S2_R4S2 | Averge | S2_R4S1 | S2_R4S2 | Averge | S2_R4S1 | S2_R4S2 | Averge |
| M_CULT_DOM-SYL_ | | 0,25 | 0,25 | 0,25 | - | - | - | 0,02 | 0,04 | 0,03 | - | - | - | 0,02 | 0,47 | 0,25 | - | - | - | 0,02 | 0,07 | 0,05 |
| M_SYL-CULT_DOM_ | | 0,12 | 0,11 | 0,12 | - | - | - | 0,01 | 0,02 | 0,02 | - | - | - | 0,01 | 0,43 | 0,22 | - | - | - | 0,01 | 0,02 | 0,02 |
| M_SYL-CULT_ROOT_EU_ | | 0,26 | 0,25 | 0,26 | - | - | - | 0,01 | 0,01 | 0,01 | - | - | - | 0,01 | 0,49 | 0,25 | - | - | - | 0,01 | 0,06 | 0,04 |
| M_CULT_ROOT_CH-BAC_HUP_ | | 0,25 | 0,23 | 0,24 | - | - | - | 0,01 | 0,01 | 0,01 | - | - | - | 0,01 | 0,49 | 0,25 | - | - | - | 0,01 | 0,09 | 0,05 |
| M_BAC_HUP-CULT_ROOT_CH_ | | 0,22 | 0,20 | 0,21 | - | - | - | 0,02 | 0,02 | 0,02 | - | - | - | 0,02 | 0,46 | 0,24 | - | - | - | 0,02 | 0,03 | 0,02 |
| M_BAC_HUP-CULT_CHN_ | | 0,19 | 0,20 | 0,19 | - | - | - | 0,04 | 0,04 | 0,04 | - | - | - | 0,04 | 0,46 | 0,25 | - | - | - | 0,04 | 0,02 | 0,03 |
| M_BAC_HUP-CULT_ROOT_EU_ | | 0,11 | 0,10 | 0,11 | - | - | - | 0,01 | 0,01 | 0,01 | - | - | - | 0,01 | 0,30 | 0,16 | - | - | - | 0,01 | 0,04 | 0,03 |
| M_CULT_CHN-BAC_HUP_ | | 0,18 | 0,27 | 0,23 | - | - | - | 0,01 | 0,02 | 0,01 | - | - | - | 0,01 | 0,49 | 0,25 | - | - | - | 0,01 | 0,10 | 0,05 |
| M_CULT_ROOT_EU-SYL_ | | 0,08 | 0,08 | 0,08 | - | - | - | 0,00 | 0,00 | 0,00 | - | - | - | 0,00 | 0,41 | 0,20 | - | - | - | 0,00 | 0,03 | 0,02 |
| M_CULT_ROOT_EU-BAC_HUP_ | | 0,16 | 0,11 | 0,13 | - | - | - | 0,01 | 0,01 | 0,01 | - | - | - | 0,01 | 0,47 | 0,24 | - | - | - | 0,01 | 0,03 | 0,02 |
| N_ANC_ | | 56115 | 57363 | 56739 | - | - | - | 8500 | 8862 | 8681 | - | - | - | 95974 | 95974 | 95974 | - | - | - | 0,07 | 0,10 | 0,09 |
| N_BAC_HUP_ | | 15372 | 19212 | 17292 | - | - | - | 91 | 67 | 79 | - | - | - | 49139 | 49139 | 49139 | - | - | - | 0,52 | 0,56 | 0,54 |
| N_CULT_CHN_ | | 10451 | 9589 | 10020 | - | - | - | 70 | 67 | 69 | - | - | - | 43875 | 43875 | 43875 | - | - | - | 0,51 | 0,59 | 0,55 |
| N_CULT_DOM_ | | 16156 | 11031 | 13594 | - | - | - | 182 | 118 | 150 | - | - | - | 48758 | 43107 | 45933 | - | - | - | 0,36 | 0,33 | 0,35 |
| N_CULT_ROOT_CN_ | | 4342 | 3106 | 3724 | - | - | - | 52 | 58 | 55 | - | - | - | 34762 | 34762 | 34762 | - | - | - | 0,48 | 0,52 | 0,50 |
| N_CULT_ROOT_EU_ | | 13423 | 7552 | 10487 | - | - | - | 68 | 51 | 60 | - | - | - | 46020 | 43494 | 44757 | - | - | - | 0,48 | 0,63 | 0,56 |
| N_SIE_ | | 20503 | 25967 | 23235 | - | - | - | 3004 | 3736 | 3370 | - | - | - | 46125 | 48073 | 47099 | - | - | - | 0,05 | 0,05 | 0,05 |
| N_SYL_ | | 9603 | 6801 | 8202 | - | - | - | 50 | 51 | 51 | - | - | - | 36011 | 31107 | 33559 | - | - | - | 0,59 | 0,59 | 0,59 |
| T_CULT_CHN-SIE_ | T_CULT_CHN-CULT_DOM_ | 176 | 143 | 160 | 1762 | 1433 | 1597 | 11 | 11 | 11 | 110 | 110 | 110 | 644 | 515 | 580 | 6440 | 5150 | 5795 | 0,12 | 0,07 | 0,09 |
| T_CULT_DOM-SIE_ | | 390 | 360 | 375 | 3896 | 3601 | 3748 | 20 | 42 | 31 | 200 | 419 | 309 | 977 | 934 | 956 | 9770 | 9340 | 9555 | 0,09 | 0,07 | 0,08 |
| T_CULT_ROOT_CN-CULT_DOM_ | T_CULT_ROOT_CN-SIE_ | 104 | 316 | 210 | 1037 | 3163 | 2100 | 10 | 11 | 11 | 100 | 110 | 105 | 328 | 101 | 215 | 3280 | 1010 | 2145 | 0,08 | 0,15 | 0,11 |
| T_CULT_ROOT_EU-SIE_ | | 257 | 274 | 265 | 2565 | 2737 | 2651 | 11 | 12 | 12 | 110 | 120 | 115 | 910 | 193 | 552 | 9100 | 1930 | 5515 | 0,14 | 0,14 | 0,14 |
| T_SIE-ANC_ | | 47645 | 50517 | 49081 | 476448 | 505174 | 490811 | 1257 | 1819 | 1538 | 12570 | 18190 | 15380 | 350278 | 327286 | 338782 | 3502780 | 3272860 | 3387820 | 0,27 | 0,25 | 0,26 |
| T_SYL-ANC_ | | 156995 | 167318 | 162156 | 1569947 | 1673177 | 1621562 | 8173 | 6611 | 7392 | 81730 | 66110 | 73920 | 470364 | 573499 | 521932 | 4703640 | 5734990 | 5219315 | 0,11 | 0,13 | 0,12 |
| T_BAC_HUP-ANC_ | | 304145 | 374483 | 339314 | 3041451 | 3744830 | 3393141 | 54905 | 53859 | 54382 | 549050 | 538590 | 543820 | 786944 | 1022054 | 904499 | 7869440 | 10220540 | 9044990 | 0,04 | 0,05 | 0,05 |

Note: NMAE, normalized mean absolute error; Divergence time were multiplied by the 10 years generation time estimates to transform them in years.

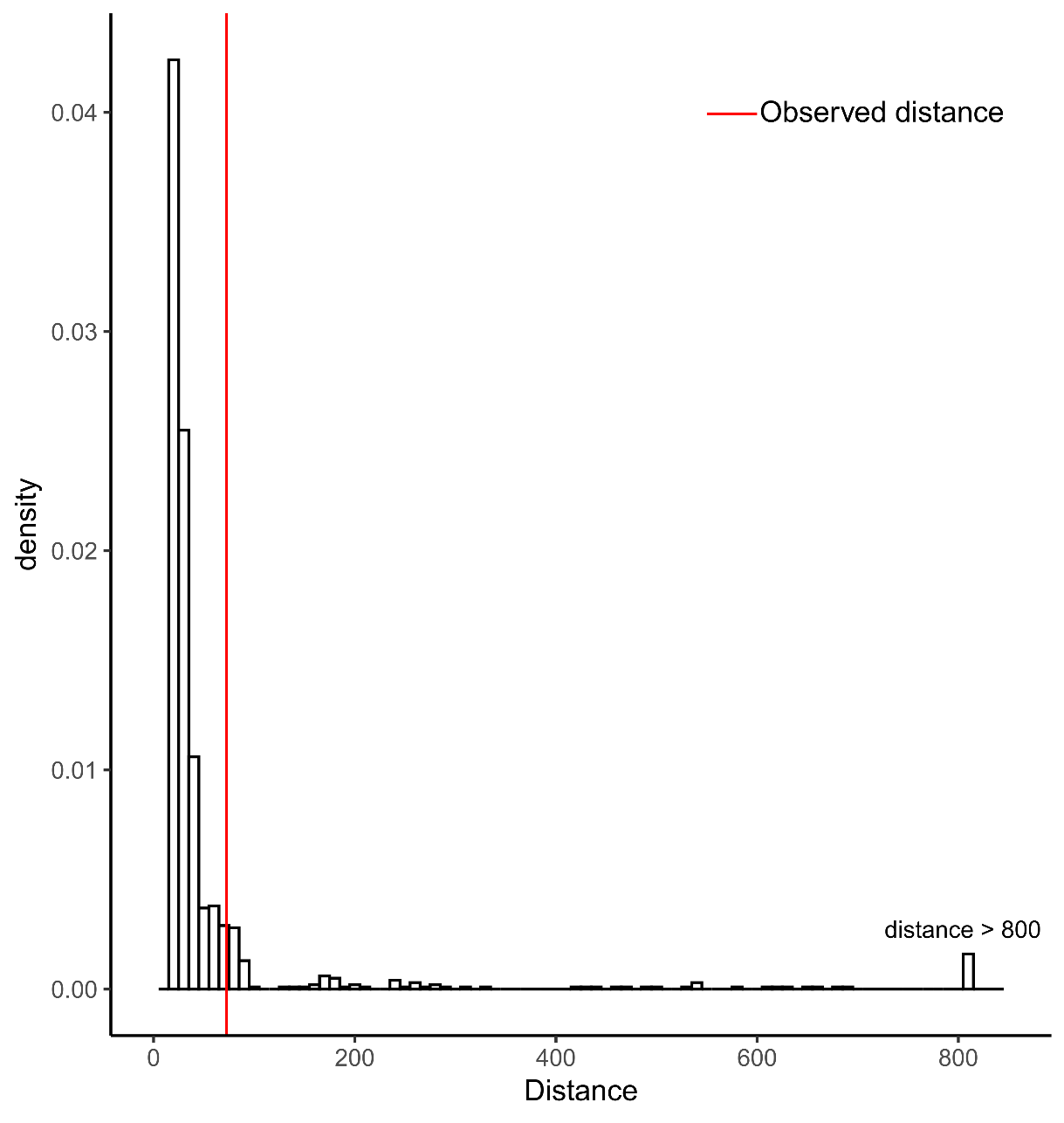

**Figure S15.** **Histogram of the 1,000 simulations of the most likely scenario of the apple domestication history inferred with ABC (*i.e*., S2_R4S2, Figure 3).** The pseudo-observed dataset was obtained using prior distributions drawn from the 90% confidence interval of the parameters estimated previously for S2_R4S2 (Table S13). This histogram is plotted under H0 (*i.e.*, the simulated dataset fits with the observed dataset) and results were obtained with the goodness of fit test from abc R package (Csilléry *et al.*, 2012), (*P*=0.069) was non-significant further confirming our model choice.

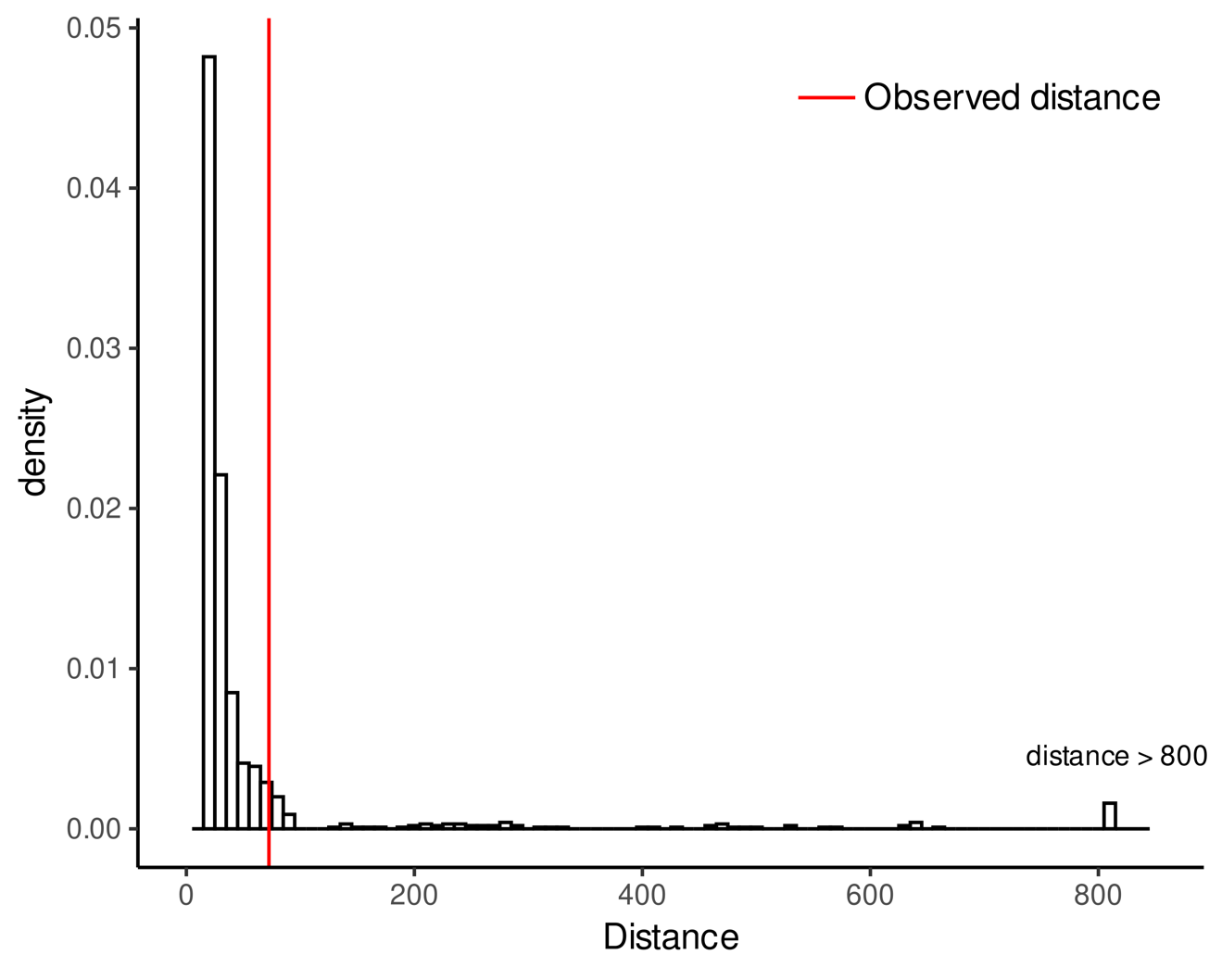
**Figure S16.** **Histogram of the 1000 simulations of the most likely scenario of the apple domestication history inferred with ABC (*i.e*., S2_R4S1, Figure 3).** The pseudo-observed dataset was obtained using prior distributions drawn from the 90% confidence interval of the parameters estimated previously for S2_R4S2 (Table S13). This histogram is plotted under H0 (*i.e.*, the simulated dataset fits with the observed dataset) and results were obtained with the goodness of fit test from abc R package (Csilléry *et al.*, 2012), (*P*=0.074) was non-significant further confirming our model choice.
